# Heatwave winners and losers: cryptic coral holobionts differ in thermal tolerance

**DOI:** 10.64898/2026.04.02.716210

**Authors:** Zoe Meziere, Ilha Byrne, Iva Popovic, Andrew Khalil, Adriana Humanes, James Guest, Cheong Xin Chan, Cynthia Riginos, Katrina McGuigan

## Abstract

Extreme climatic events are reshaping ecosystems worldwide as individual organisms vary markedly in their ability to withstand these disturbances. Deciphering patterns of persistence on local scales is therefore critical for predicting biodiversity trajectories under intensifying climate extremes. In this study, we examined variation in thermal stress responses among individuals of the coral *Stylophora pistillata* species complex during a heatwave at Heron Island Reef, Australia. More than half of the focal coral colonies died on the reef, and survival of coral fragments maintained under *ex situ* common thermal stress conditions was significantly correlated with the survival of their source colony. This demonstrates that survival differences result largely from biological factors rather than differential thermal exposure across reef habitats. Under common garden conditions, we observed striking differences in bleaching severity and survival times among three sympatric cryptic taxa and their highly host-specific symbiont community. Within the most locally common taxon, corals from historically warmer and more seasonally variable reef habitats seem more susceptible to bleaching, contrary to expectations. Together, these results reveal how biological differences among cryptic taxa and among individuals can shape coral responses during a heatwave and advance our understanding of coral vulnerability in a rapidly warming world.

## 1. Introduction

Why do some individuals survive extreme climatic events while others do not? Organisms at a given location are rarely affected equally by acute environmental perturbations such as droughts, heatwaves or floods. Understanding the factors driving this variation has never been more urgent given the increasing frequency, intensity, and duration of climate anomalies (Easterling et al., 2000; Ummenhofer & Meehl, 2017). To assess vulnerability to an extreme event, we need to differentiate between the factors that determine sensitivity and the factors that govern exposure (Turner et al., 2003). In this framework, sensitivity is driven by traits that are intrinsic to a species whereas exposure reflects extrinsic factors determined by local habitat effects (Williams et al., 2008).

Species inhabiting the same ecosystem often have different sensitivity to a given environmental stress, resulting in different survival outcomes following an extreme event. Large-body, short-lived, low-dispersal and highly specialised species, for example, have been theorised to be more susceptible (Pimm et al., 1988). While tolerant species will tend to survive, more sensitive species might exhibit higher risks of local population decline or extinction (Thomas et al., 2004). As a result, climate extremes are rapidly altering species richness and community compositions across many ecosystems (Ummenhofer & Meehl, 2017). At the same time, most ecosystems are structurally complex with different (micro)habitats, and this fine-scale environmental variation can buffer organisms from extreme conditions (reviewed in Denney et al., 2020). For instance, leaf litter and tree holes can regulate temperature and moisture regimes within a rainforest and have been shown to become local thermal refugia during hot periods (Scheffers et al., 2014). As a result, individuals inhabiting adjacent habitats might experience different exposure to a given climatic event and consequently exhibit different phenotypic responses. Therefore, survival under climate extremes can depend on both biological and environmental factors. The rarity and unpredictability of extreme climate events present a major challenge to disentangling these factors (Altwegg et al., 2017), yet this information is important for predicting and managing biodiversity under intensifying climate change.

Tropical coral reefs provide a striking example of ecosystems increasingly threatened by extreme climatic events, particularly marine heatwaves (Hughes et al., 2017; Sully et al., 2019). Marine heatwaves can last several weeks and significant variation in thermal stress responses are often observed among coral colonies inhabiting the same reef (e.g., Hoogenboom et al., 2017; Hughes et al., 2017). Intrinsically, not all coral species exhibit the same sensitivity to thermal stress. For example, a coral species’ morphotype (e.g., Loya et al., 2001; van Woesik et al., 2011), tissue thickness (e.g., Loya et al., 2001), or ability to feed heterotrophically (e.g., Conti-Jerpe et al., 2020, 2020; Rodrigues & Grottoli, 2007) may impact its thermal sensitivity. Although morphologically similar species are often assumed to be ecologically equivalent (Bickford et al., 2007), growing evidence indicates that morphologically cryptic coral species can also exhibit different bleaching responses (e.g., Burgess et al., 2021; Starko et al., 2023).

Additionally, the coral-algal symbiosis is an important intrinsic factor shaping holobiont fitness. Coral growth and calcification largely relies on the translocation of compounds from photosynthetic dinoflagellates of the family *Symbiodiniaceae* (Gattuso et al., 1999). Coral species differ in how this partnership is established: while some acquire symbionts from the environment, others transmit them maternally, a mechanism that is expected to result in greater symbiont-host specificity (Baker, 2003). Similarly, host coral species can form strict associations with only one *Symbiodiniaceae* taxon or associate with multiple taxa. This variation in symbiont acquisition matters because some symbiont taxa confer greater heat tolerance to the holobiont than others (e.g., Berkelmans & van Oppen, 2006; Rouzé et al., 2019), and a more flexible symbiosis may allow for the acquisition of more heat-tolerant symbionts during heat stress (Fabina et al., 2012). When symbiosis is disrupted, the resulting loss of photosynthetic symbionts leads to coral bleaching and ultimately death if symbiosis is not restored (LaJeunesse et al., 2018). However, symbiont community composition within a coral colony cannot be assessed through visual surveys and the coral microbiome therefore represents another hidden player that could explain differential coral survival during thermal stress.

Importantly, heatwaves do not uniformly affect all parts of the reef and thermal stress exposure therefore varies across habitats. For example, reef flats and lagoons are subject to strong tidal oscillations and can reach very high temperatures on low tides. While locations exposed to currents and wave action typically remain cooler, neighbouring locations sheltered from constant water movement are warmer (Brown et al., 2023; Green et al., 2019). As a result, some reef habitats might experience disproportionate levels of heat exposure during a heatwave, which could also explain why corals inhabiting these habitats are more affected. Notably, variation in thermal exposure across habitats can confound observations of species-specific responses during a heatwave if the species of interest occupy different habitats. In particular, this might be a challenge when studying thermal sensitivity differences in coral species complexes, as cryptic species often inhabit distinct environments (reviewed in Grupstra et al., 2024).

In addition to intrinsic differences in thermal sensitivity among species, conspecific individuals occupying environmentally distinct habitats might also differ in environmental sensitivity if they are locally acclimatised or adapted. While long-term acclimatisation can induce persistent physiological adjustments in response to the local environmental conditions, natural selection acts over multiple generations to favour genotypes with optimal phenotypes (Chevin et al., 2010; Kawecki & Ebert, 2004). Both processes result in phenotypes (either plastic or genetic) that match their environment and could therefore explain variation in intra-specific responses among habitats during an extreme climatic event (Reed et al., 2011; Somero, 2010). Previous studies have found evidence for fine-scale thermal sensitivity differences among conspecific coral colonies under experimental heat stress (e.g., Humanes et al., 2022; Naugle et al., 2024). However, fewer studies have related this thermal sensitivity variation to habitat-specific historical thermal regimes, so it remains unclear whether it is adaptive (but see Oliver & Palumbi, 2011; Schoepf et al., 2015; Thomas et al., 2022).

Advancing our understanding of variation in coral thermal stress responses requires separating intrinsic factors affecting sensitivity (i.e., identity and genotype of the host and the symbionts) from the extrinsic factor of thermal stress exposure. Common garden experiments have long been used to test intrinsic effects, by growing or keeping individuals originating from different habitats in the same environment (Berend et al., 2019; de Villemereuil et al., 2016). To further clarify the effect of environmental history on the phenotypes expressed under common conditions, we can determine how much of the phenotypic variation can be attributed to environmental variation among the natal habitats (Blanquart et al., 2013; Hereford, 2009). Although coral research has extensively utilised experimental designs where coral fragments are all exposed to the same thermal stress, these experiments have not typically applied heat stress regimes that reflect realistic heatwave conditions (McLachlan et al., 2020), have not considered variation in recovery trajectories following the heat stress, and it remains unclear whether *ex situ* responses of fragments reflect whole-colony responses on the reef (Grottoli et al., 2021).

In this study, we took advantage of a real-time marine heatwave to investigate thermal sensitivity variation among *Stylophora pistillata* corals, integrating *in situ* observations with *ex situ* common garden experiment. *Stylophora pistillata* is a brooding coral with meter-scale larval dispersal (Meziere et al., 2025), an early coloniser following disturbances (Loya, 1976), and was recently identified as a complex of cryptic and sympatric species across the Great Barrier Reef (Meziere et al., 2024). *Stylophora pistillata* is categorised among the most thermally susceptible corals (e.g., Loya et al., 2001; Marshall & Baird, 2000), however, previous studies did not account for the existence of cryptic taxa. Other studies have also documented high intra-specific bleaching variation among *S. pistillata* colonies in the field (e.g., Sampayo et al., 2008), consistent with cryptic species that differ in thermal stress sensitivity. These attributes make *S. pistillata* an ecologically interesting and tractable system to investigate response variation within and among species under thermal stress. Our goals were to (1) quantify thermal stress response variation among coral colonies during a marine heatwave, (2) test the importance of intrinsic sensitivity differences between cryptic coral holobionts, and (3) compare coral colony survival under common garden and *in situ* to assess the effect of habitat-specific thermal stress exposure. We predicted both among-species and within-species thermal stress response variation, reflecting differences in intrinsic sensitivity and local adaptation to contrasting thermal habitats.

## 2. Methods

### 2.1. Bleaching event, coral collection and common garden experiment

In 2024, a mass coral bleaching affected the Southern Great Barrier Reef (GBR) of Australia (Henley et al., 2024). At Heron Island Reef, the heatwave lasted from mid-January to May and resulted in coral bleaching across 65% of the reef slope (Rowell et al., 2026). Shortly after the onset of the heatwave, 3-7 February inclusive, we sampled 80 *S. pistillata* colonies from ten locations (4-12 colonies per location) across Heron Island Reef (Table S1, Table S2) following depth transects from 12 meters to 2 meters (Marine Parks Permit number G21/44774.1). Each coral was photographed and tagged, and we collected three 5 cm length fragments (used in the common gardens, n = 240) using a hammer and a chisel and one 2 cm^2^ tissue sample (used for DNA sequencing, n=80) using pliers. Within two hours after collection, the fragments were glued onto labelled aragonite substrata and placed in aquaria (Figure S1). Tissue samples were fixed in absolute ethanol and then placed in a −20*°*C freezer. Sampled colonies were revisited 2.5 months after initial collections (15-19 April 2024 inclusive), when temperatures had returned below historical maximum monthly mean, to assess their survival.

Common garden exposure lasted 84 days (3 February-26 April), largely spanning the heatwave duration. Seawater was sourced at 15 meters depth between Heron Island Reef and Wistari Reef and was pre-filtered to exclude particles larger than 3 mm. We created a semi flow-through system with two outdoor tables, three sumps per table and two 70L aquarium tanks per sump (Supplementary Material, Figure S1). Water temperature was not manipulated and followed natural diel fluctuations (Figure 1D) with similar temperature profiles among tanks (mean sd = 0.1°C; Figure S2). On coral reefs, the intensity and duration of heat stress is commonly quantified in Degree Heating Week (DHW; °C-wk). We calculated DHW prior to the start of the common garden experiment and for the duration of the experiment (See Supplementary Material).

**Figure 1.**
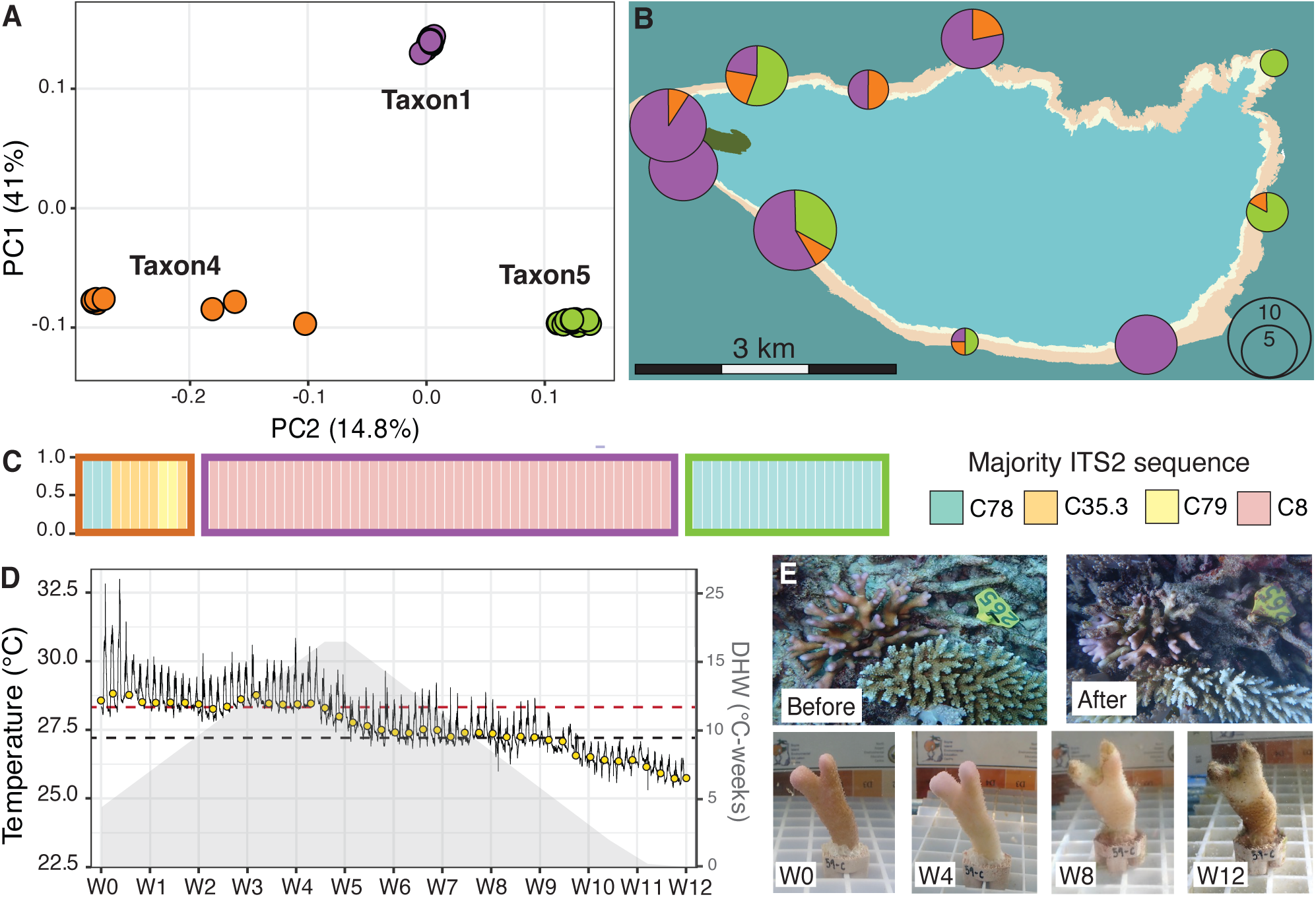
Coral holobiont identities of *Stylophora pistillata* corals at Heron Island Reef and common garden conditions during a heatwave. (A) Principal Component Analysis reveals three taxa, previously identified as *S. pistillata* Taxon1, Taxon4 and Taxon5. PC1 and PC2 axes reversed for visualisation; (B) Map showing the distribution and sampling effort for all taxa; (C) ITS2 relative abundance for each coral colony, coloured by majority ITS2 sequence. Taxon4 samples are ordered to match host differentiation along PC2; (D) Temperature profile in the common garden tanks (average aquarium loggers, black solid line) and on the reef (NOAA CRW 5 km, yellow dots). The black dashed line represents the climatic maximum monthly mean (MMM), the red dashed line represents the bleaching threshold (1°C above MMM), the grey area represents the accumulation of thermal stress over time; (E) photographs of a *S. pistillata* coral colony *in situ* before and after the heatwave (upper left coral colony on the image), and a fragment of the same colony in the common garden between week 0 (W0) and week 12 (W12).

### 2.2. Coral host taxa delineation

We extracted genomic DNA using the DNAeasy QIAGEN Blood and Tissue kit according to the manufacturer’s instructions and prepared whole-genome libraries using the Lotus DNA Library Prep Kit for NGS with 10 ng of input DNA and enzymatic fragmentation to achieve average insert sizes of 350 bp. Libraries were sequenced on a single shared NovaSeq 6000 S4 flowcell late (300 cycles, 150 bp paired-end reads). We used Trimmomatic (Bolger et al., 2014) to quality-filter reads, BWA-MEM (Li & Durbin, 2009) to map reads to the *S. pistillata* reference genome (GCA_032172095.1), Samtools v1.10 (Danecek et al., 2021) to index BAM files and picard (http://broadinstitute.github.io/picard/) to remove PCR duplicates. We used GATK call variants (Gamer et al., 2019), VCFtools v0.1.16 (Danecek et al., 2011) to filter the dataset and PLINK v2.0 (www.cog-genomics.org/plink/2.0/, Chang et al., 2015) to perform identity-by-state and PCA analyses to identify clones and assess genetic structure, respectively. Details on this pipeline can be found in Supplementary Material.

### 2.3. *Symbiodiniaceae* community composition assessment

We assessed the community of symbiotic dinoflagellates (family *Symbiodiniaceae*) for each sampled coral colony (n=80) and each fragment that survived the thermal stress (n=76) using Internal Transcribed Spacer 2 (ITS2) profiling. Amplicon-based sequencing was performed at the Ramacciotti Centre for Genomics on a NextSeq 1000 P1 platform with 2x300bp using the *Symbiodiniaceae*-specific primers (Hume et al., 2013), and a PCR duplicate for each sample (total of 312 samples). Duplicates were sequenced and analysed separately to evaluate variation between PCR reactions. *Symbiodiniaceae* species can be difficult to delineate because of high ITS2 intragenomic sequence diversity (Davies et al., 2023). We therefore used SymPortal (Hume et al., 2019), which assigns ITS2 type profiles based on sequence variants that co-occur in multiple samples and are likely to belong to the same species. Six samples did not yield any sequence data and were therefore not submitted to SymPortal. For the remaining 606 samples, eight samples had low read counts (< 7,000 sequence variant reads and/or < 6,000 ITS2 profile reads; Supplementary Material) and were discarded from downstream analyses. All duplicates had the same ITS2 profile (Bray–Curtis similarity of profile relative abundance 0.98-1.00), confirming no PCR artefacts (Figure S3), and we kept one sample per duplicate pair.

ITS2 sequence variant composition was analysed using non-metric multidimensional scaling (NMDS). To reduce the influence of rare and highly abundant sequences, we standardized the count data using Wisconsin double standardization, before calculating pairwise Bray-Curtis dissimilarities among samples and running the NMDS using the ‘vegan’ R package (Oksanen et al., 2015). Then, to test the effect of the host taxonomic identity, sampling location, sampling depth, and experimental exposure (i.e., wild colony vs. survivor experimental fragment) on ITS2 composition, we used permutations to generate a null distribution of the data under no effects of the predictors. This was performed using PERMANOVA (999 permutations) implemented in the ‘vegan’ R package. To account for non-independence among samples of the same coral colony (e.g., common garden fragments and wild fragments), permutations were restricted within host genotypes, such that replicates were permuted within colonies and not exchanged among colonies.

### 2.4. Historical temperature characterization at sampling sites

While detailed environmental data for each sampled coral colony would have been ideal, obtaining such information was not feasible in this study. Instead, we used modelled data from the GBR1 hydrodynamic model database (Steven et al., 2019), using the geographic coordinates of the sampling sites and the sampling depth for each coral colony to extract data. We used the monthly products and the longest date range available (01/12/2014 – 01/01/2024), to capture long-term environmental effects, and the following eReefs depth bins: 0.5-2.5m, 2.5-5.25m, 5.25-9m, 9-13m. From the raw temperature data, we calculated four ecologically meaningful variables: mean annual temperature (captures baseline thermal conditions), minimum monthly mean temperature (captures winter cold), maximum monthly mean temperature (captures summer heat) and annual temperature range (captures seasonal variability). These parameters varied only slightly among sites (Table S1) but these small differences may be biologically meaningful. These variables were highly correlated (Pearson’s r > 0.7), so we performed a PCA and retained the first two axes (temp-PC1, and temp-PC2), which captured most of the variation in the thermal environment (91% and 8%, respectively) (Figure S5). Temp-PC1 strongly associated with longitude (east vs west) and captured thermal variability (high values reflecting with warmer temperature maximum, colder temperature minimum and greater temperature range). Temp-PC2 strongly associated with latitude (leeward vs windward) and captured mean temperature (high values reflecting lower mean temperature).

### 2.5. Fragment phenotypic measurements

Every seven days, fragment survival, bleaching status, bleaching percentage and health score were assessed. Survival was classified as alive (entire fragment covered in live tissue), partially dead (part of the fragment without live tissue) or dead (no live tissue visible). Bleaching status was categorised as no bleaching (no tissue discolouration), partial bleaching (part of the fragment with tissue discolouration) or full bleaching (fragment fully discoloured). Bleaching area was visually estimated and standardised by fragment total surface area, ranging from 0% (no bleaching) to 100% (full bleaching) in 5% increments. Health score was determined using the CoralWatch Health Chart and ranged from 1 (very pale) to 6 (very dark) (Siebeck et al., 2006). Visual scores should not be interpreted as exact values, but as qualitative categories allowing consistent comparisons across fragments and timepoints.

To capture important aspects of phenotypic responses over time, we calculated four phenotypic response variables: (1) survival time (i.e., the number of weeks a fragment stayed alive, proxy for persistence); (2) time without bleaching, (i.e., the number of weeks between the start of the experiment and the first visual signs of bleaching, proxy for bleaching resistance); (3) maximum bleaching area per fragment (proxy for bleaching severity); and (4) recovery time (i.e., number of weeks between a fragment’s lowest health score and the first subsequent increase in its health score, proxy for resilience). For (1), (2) and (4), we used a census value of 12 weeks for fragments that (1) survived until the termination of the experiment, (2) never showed signs of bleaching and (3) did not improve their health score,. Only fragments that were alive by the end of the experiment and showed signs of paling were assigned a recovery time value.

### 2.6. Survival and bleaching response differences among taxa

All statistical analyses were conducted in R v. 3.5.0 (R Core Team, 2023). First, differences in survival probability over time among taxa were assessed using a mixed-effects Cox model in the ‘coxme’ R package (Therneau, 2009). We also ran this analysis to investigate survival and bleaching probability over DHW exposure (see Supplementary Material). We visualised survival trajectories with Kaplan–Meier survival curves using the ‘survival’ R package (Therneau, 2001). Second, we fitted Generalised Mixed Linear Models (GLMMs) using the ‘glmmTMB’ R package (McGillycuddy et al., 2025) for the four traits described above: survival time, time without bleaching, maximum bleaching area and recovery time. In both survival models and GLMMs, we used ‘Taxon’ as a fixed effect, ‘Tank’ (nested within ‘Sump’) and ‘Colony’ (nested within ‘Taxon’) as random effects to account for uncontrolled environmental effects among tanks and colony-level effects, respectively. Any remaining, unexplained, variation among replicate fragments of the same colony are estimated as the model residual (see details in Supplementary Material). For the GLMMs, significance of fixed effects was assessed using likelihood ratio tests implemented the ‘car’ R package (Weisberg & Fox, 2011) and estimated marginal means with Tukey post hoc pairwise comparisons were obtained using the ‘emmeans’ R package (Lenth, 2017).

### 2.7. Survival and bleaching response differences within taxon among environments

To test whether conspecifics originating from different thermal environments (Figure S5) showed different responses, we focused on variation among Taxon1 colonies, since it was the most sampled taxon (see Results). We fitted GLMMs for the four phenotypic traits described above, but to capture variation explained by the historical thermal environment, we used temp-PC1 and temp-PC2 as the fixed effects (see Supplementary Material). We excluded three sampling sites (BA1, FR1 and Libby’s Lair) with very low sample size (< 5 Taxon1 colonies), to limit the potential for unrepresentative samples to bias interpretations. We also omitted two locations (H5A and H5B) where Taxon1 was not sampled. Random effects were again fit to account for data structure as detailed above.

### 2.8. Survival differences under different heat stress exposures on the reef

If corals on the reef experienced different thermal exposures during the heatwave, then survival under the *ex situ* common garden conditions should be a poor predictor of *in situ* survival. To assess this, we estimated Cohen’s Kappa (treating common garden vs. *in situ* as independent raters of survival) and tested the null hypothesis of no agreement (Kappa = 0) using the ‘irr’ R package (Gamer et al., 2019). Analyses were conducted for all colonies together, and for each taxon separately. Only colonies that could be re-assessed post-heatwave were included. For the common garden data, we assigned a colony-level score to account for the three replicate fragments, using “alive” if at least two out of three fragments were alive at the last time-point and “dead” if two or three fragments were dead. The common garden experiment was primarily designed to expose all fragments to a standardized, yet realistic, heatwave thermal regime. While similar, the thermal conditions in the common garden are unlikely to exactly replicate the conditions experienced by the colonies on the reef. This is a limitation precluding conclusions about differences in average survival ex situ versus in situ but does not impact conclusions about whether survival was an intrinsic response of the corals (i.e., whether colonies with higher ex situ survival also had higher survival *in situ*).

## 3. Results

### 3.1. Genomic data processing and filtering and host taxa delineation

Whole-genome sequencing of the 80 *S. pistillata* corals yielded a mean read coverage of 10.2x. After variant filtering, we retained 2,021,787 SNPs. For PCA, we removed linked sites, resulting in 712,815 SNPs. No clones were detected. PCA of genomic data showed three distinct clusters (Figure 1A). Meziere et al., (2024) discovered four divergent taxa within *S. pistillata* across the GBR (labelled Taxon1, Taxon3, Taxon4 and Taxon5). In the present study, we re-sequenced four representative individuals of Meziere et al., (2024) for each of the four taxa. Including these representative individuals in our dataset, we assigned the colonies sampled in this study to *S. pistillata* Taxon1 (n=40), Taxon4 (n=11) and Taxon5 (n=20) (Figure S6A). Three samples within our dataset appeared taxonomically intermediate (Figure 1A), which was not consistent with levels of missing data (Figure S6B). Downsampling to five samples per taxon resulted in more cohesive grouping of these samples with Taxon4 representative samples (Figure S6C), suggesting that their intermediate PCA position could be an artefact of uneven sample numbers among genetic clusters. Nonetheless, ADMIXTURE analysis showed that one of these individuals might be of hybrid origin (Figure S7). We assigned this sample to Taxon4, due to its position on the PCA and the greater Taxon4 ancestry proportion, but we also conducted all statistical analyses after excluding this sample to confirm that results were consistent. Notably, we sampled at least two taxa at most (7/10) sites (Figure 1B) with no clear partitioning of taxa across water depth (Figure S8).

### 3.2. Symbiodiniaceae community composition assessment

We identified four ITS2 type profiles associated with the 80 original colonies, all belonging to *Symbiodiniaceae* Clade C: C8/C8a-C1-C42.2-C42e-C8e-C42ae (n=49 Taxon1 coral hosts), C78/C78c-C1-C3lk-C3gk-C1re-C78d (n=2 Taxon4 and n=21 Taxon5 coral hosts), C35.3/C35.1/C35a-C54l (n=6 Taxon4 coral hosts), and C79/C54l-C79b-C79a-C40ap-C54m (n=2 Taxon4 coral hosts) (Figure 1C, Figure S3, Figure S4). In the next sections, we refer to these type profiles using the majority ITS2 sequence (i.e., C8, C78, C35.3 and C79). Each coral was dominated by one ITS2 type profile (i.e., not a mixed community), and, with the exception of C78, ITS2 profiles were highly specific to the host taxon. Intriguingly, the only three surviving fragments of Taxon4 were all from the same colony (colony #66), which harboured C78 symbionts, and this colony was also the only Taxon4 colony found alive on the reef post-heatwave. We also performed ITS2 sequencing for the 76 surviving fragments at the end of the experiment to compare symbiont communities pre- and post- heat stress. Only four fragments exhibited ITS2 profiles different from those before the heatwave: the three fragments of Taxon1 colony #49 switched from C8 to C78, and one fragment of Taxon1 colony #59 switched from C8 to a mixed community of C8 and A1er (Figure S3). These fragments never showed signs of bleaching.

NMDS ordination of Bray–Curtis dissimilarities revealed clear structuring of *Symbiodiniaceae* ITS2 sequence variants among samples, clustering according to the ITS2 type profiles inferred from SymPortal (Figure S4). Within clusters defined by type profiles, we found some additional differentiation according to the depth of the sampled coral colonies. PERMANOVA confirmed that ITS2 composition was significantly structured by the host taxonomic identity (*R²* = 0.184, *p* = 0.006), while the sampling location (*R²* = 0.063, *p* = 0.307), depth (*R²* = 0.005, *p* = 0.099) and experimental exposure (*R²* = 0.005, *p* = 0.008) had a comparatively weak contributions.

### 3.3. Survival and bleaching response differences among host taxa

In total, 164 out of 240 fragments (68%) were dead at the end of the experiment and 39 (16%) had partial mortality. Survival differed among taxa, with all three fragments dying in 25/49 (51%) Taxon1 colonies, in 10/11 (90%) Taxon4 colonies, and in 6/20 (30%) Taxon5 colonies (Figure 2A). Survival analyses supported taxon-specific responses with statistical significance (χ² = 15.1, *p* = 0.0005, Table S3), with lowest mortality risk in Taxon5, and highest mortality risk for Taxon4 (Figure 2B). Survival differed significantly between Taxon4 and the other taxa before peak DHW (χ² = 36.3, *p* = 1.3e-08; Figure S9A) and among all taxa after peak DHW (χ² = 14.7, *p* = 0.0006; Figure S9B), reflecting the continuing mortality of Taxon1 despite cooling water temperatures. These differences are supported by significant differences among taxa in the average number of weeks they remained alive in the common garden (χ² = 41.6, *p* < 0.001; Figure 2C, Table S3, Table S4).

**Figure 2.**
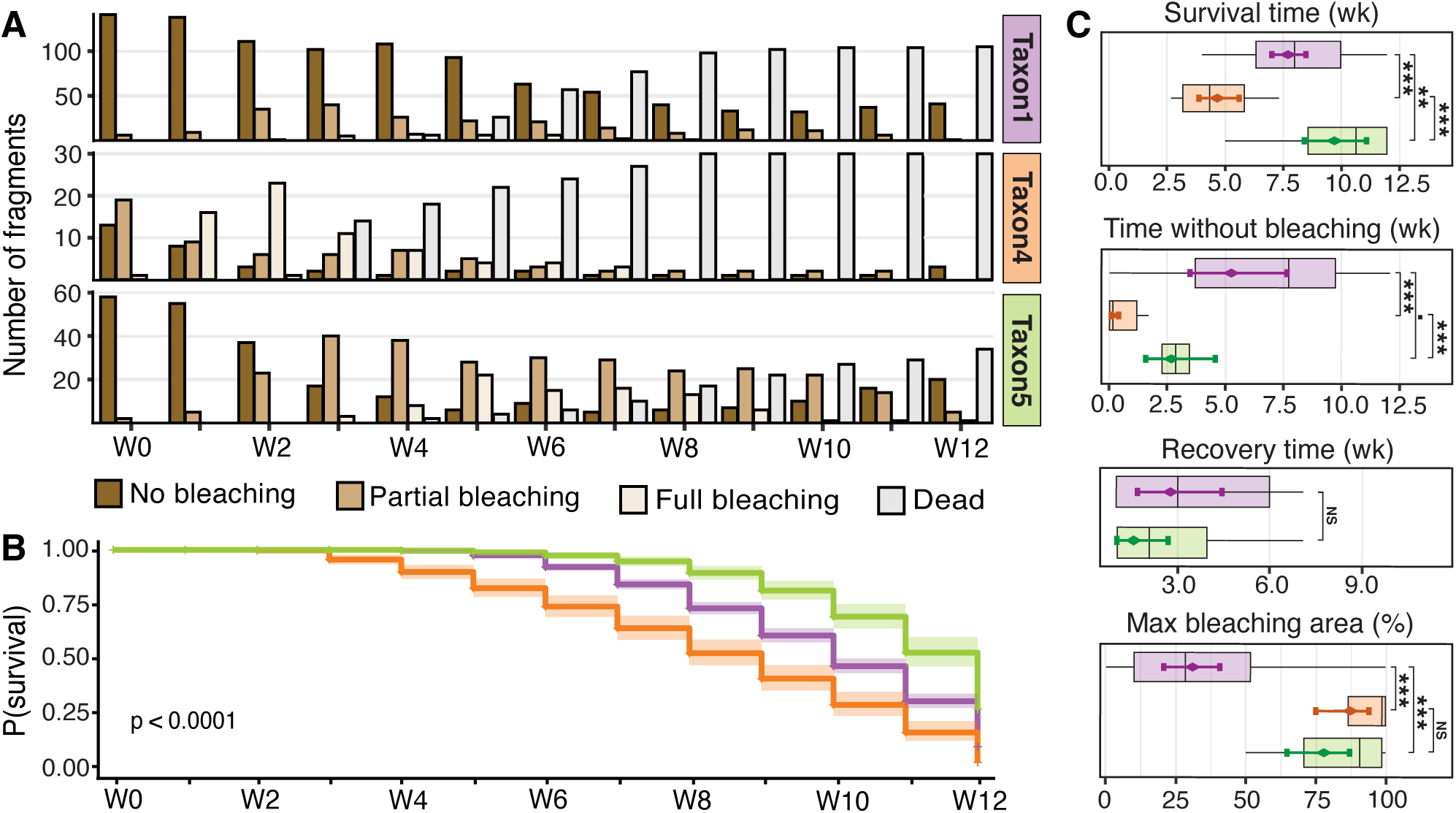
Survival and bleaching response differences among *Stylophora pistillata* taxa during twelve weeks (W0-W12) of thermal stress conditions in the common garden experiment. For each *S. pistillata* taxon, we show (A) the number of alive fragments according to their bleaching status (full, partial, none) and the number of dead fragments; (B) the probability of colony survival as Kaplan-Meier survival curves; (C) the distributions of colony-level average (across the three fragment replicates) values for: the number of weeks a colony was alive, the number of weeks a colony did not show signs of bleaching, the number of weeks it took a colony’s health score to increase after reaching its lowest value and a colony’s maximum bleached area. Recovery time for Taxon4 is not shown because fragments from only one Taxon4 colony survived. Diamonds represent model-estimated marginal means (with 95% confidence intervals) and we indicate significance of pairwise differences using: “NS” for *p* > 0.1, “*.”* for *p* < 0.1, “**\***” for *p* < 0.05, “**\*\*”** for *p* < 0.01, “**\*\*\***” for *p* < 0.001.

We observed bleaching (full or partial) in 177 (74%) fragments over the course of the experiment and bleaching recovery occurred in 46 (19%) fragments (Figure 2A). Bleaching varied significantly by taxon (χ² = 67.8, *p* < 0.001), as did the maximum bleaching area (χ² = 63.6, *p* < 0.001) (Figure 2C). Notably, while Taxon5 had the highest survival, it bleached faster and more extensively than Taxon1 (Table S3, Table S4; see also Figure S9C, D). Finally, while the recovery time was slightly faster in Taxon5 fragments, we found no significant difference in recovery time between Taxon1 and Taxon5 (Figure 2C, Table S3). We could not estimate recovery times in Taxon4 because only three fragments (from the same colony) survived until the end of the experiment. Statistical significance in survival and mixed model analyses was unaffected by the exclusion of the putative hybrid sample (see Supplementary Results).

### 3.4. Survival and bleaching response differences within taxon among environments

While temp-PC1 and temp-PC2 had no significant effect on survival time among Taxon1 colonies (temp-PC1: χ² = 1.35, *p* = 0.24; temp-PC2: χ² = 0.83, *p* = 0.36), there was weak evidence that their bleaching responses may vary along temp-PC1, which captures thermal variability and more extreme temperatures (time without bleaching: χ² = 3.21, *p* = 0.07; maximum bleaching area: χ² = 2.78, *p* = 0.09). Specifically, faster bleaching and more extensive bleaching were marginally associated with higher temperature variability (i.e., higher temp-PC1 scores). There was no evidence that bleaching responses varied along temp-PC2, which captures mean temperature (time without bleaching: χ² = 0.31, *p* = 0.57; maximum bleaching area: temp-PC2: χ² = 0.74, *p* = 0.38).

### 3.5. Survival differences under different heat stress exposures on the reef

Following the heatwave (three months after initial sampling), we located 53 (66%) of the 80 tagged source colonies. Of these 53, 12 (23%) were alive, 13 (24%) had partial mortality, and 28 (53%) were dead (Figure 3B, Figure S10). Of the 25 living colonies, 23 appeared healthy (no bleaching) and two showed partial bleaching. On-reef colony survival differed markedly among taxa (Figure 3A): nearly all Taxon4 re-located colonies (6/7) were dead, whereas 48% (15/31) of Taxon1 and 60% (9/15) of Taxon5 re-located colonies were alive, consistent with survival patterns under common garden conditions.

**Figure 3.**
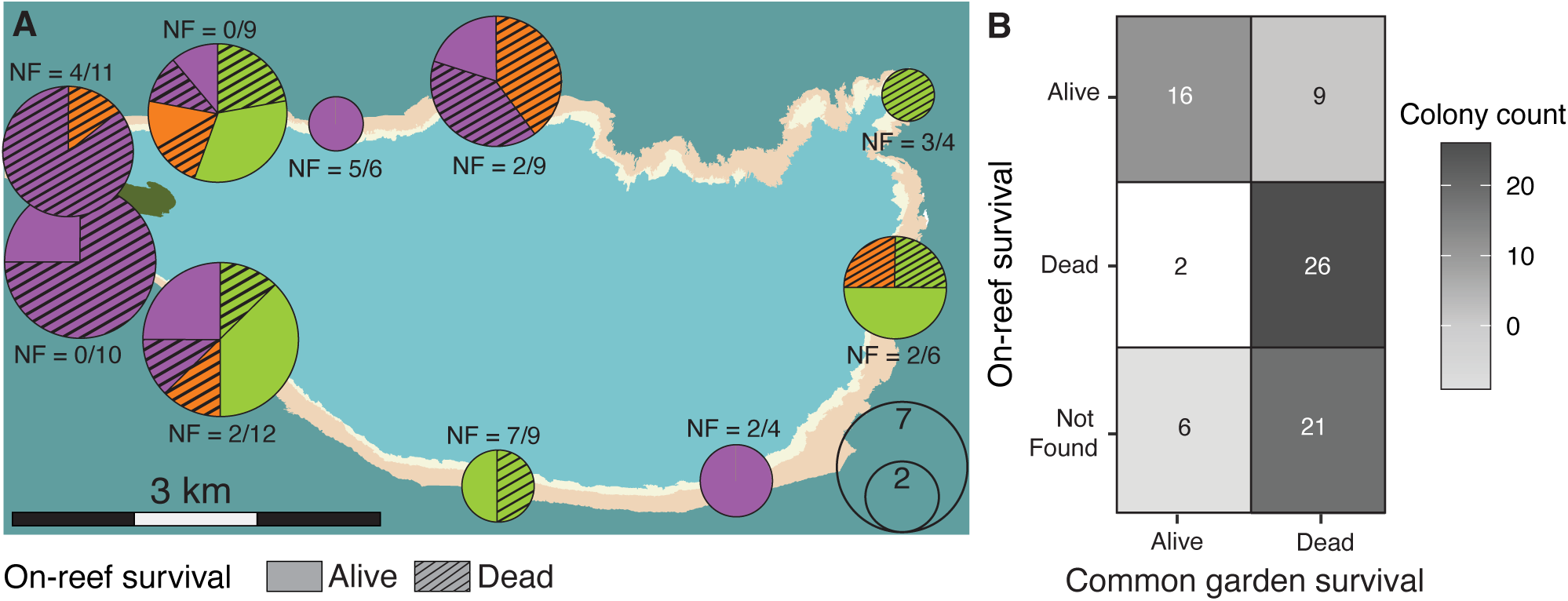
On-reef survival of tagged *Stylophora pistillata* colonies following the heatwave. (A) Map showing survival of the sampled colonies for each taxon (purple, orange and green for Taxon1, Taxon4 and Taxon5, respectively). Dead colonies are represented with stripes and alive colonies with block colours; NF indicates the number of colonies that we could not find post-heatwave; (B) heatmap showing the relationship between survival of the 80 colonies on-reef following the heatwave (rows) and in the common garden experiment (columns). Each tile shows the number of colonies for each combination (e.g., of the 25 colonies that were Alive *in situ* post-heatwave, 16 were categorised as Alive in the common garden, and 9 as Dead).

We found a significant association between colony survival under common garden conditions and on-reef survival (Kappa = 0.171, z = 4.74, *p* = 0.00018) with a Spearman’s correlation coefficient of ρ = 0.62 (Figure 3B). Interestingly, most colonies that could not be re-located on the reef had dead fragments in the common garden. Dead colonies in the field, likely overgrown with macroalgae, may have been difficult to detect, explaining our inability to find them. The association between common garden and on-reef survival was also significant within each taxon (Taxon1: Kappa = 0.121, z = 1.97, *p* = 0.049; Taxon4: Kappa = 0.462, z = 2.65, *p* = 0.008; Taxon5: Kappa = 0.130, z = 2.37, *p* = 0.018; Figure S10). This strong association of survival outcomes suggests that differential exposure to heat stress across the reef likely contributes little to differential survival.

## 4. Discussion

In this study, we explored sources of thermal tolerance variation among coral colonies of the *Stylophora pistillata* species complex during a marine heatwave at Heron Island Reef, Australia. Combining a common garden experiment and *in situ* monitoring, we found high concordance between on-reef survival and common garden survival, suggesting a negligible effect of thermal stress exposure differences among reef habitats. This finding also confirms that the common garden observations likely provided biologically meaningful insights into on-reef responses. Instead, we found most bleaching and survival differences among coral colonies reflected the intrinsic sensitivity of closely related taxa and their associated symbionts. Unexpectedly, we found some evidence for higher within-species thermal sensitivity in corals inhabiting historically warm and variable habitats.

### Cryptic taxa differ in their bleaching responses and survival

Our study was the first to pair a coral common garden experiment with *in situ* reef observations during a marine heatwave, thereby testing intrinsic differences in thermal stress response. Under common garden conditions, the three *S. pistillata* taxa showed different bleaching and survival patterns (Figure 2), demonstrating that observed inter-specific differences reflect sensitivity differences among these taxa, not differential heat stress exposure in their native habitat. Importantly, we also show that survival on the reef mirrored experimental survival (Figure 3). Across taxa, studies have revealed that previously presumed generalist species are in fact complexes of specialist cryptic species (reviewed in Bickford et al., 2007). Our findings add to this growing body of literature and lend support to previously described response diversity among cryptic coral species (e.g., Burgess et al., 2021; Rose et al., 2021; Starko et al., 2023). Variation in how cryptic species respond to environmental pressure can have far-reaching ecological consequences (Burgess et al., 2022). Under recurring climate extremes, it is plausible that more susceptible taxa (such as *S. pistillata* Taxon4) could be gradually replaced by more resilient ones (such as *S. pistillata* Taxon5). Conversely, if all cryptic species within a species complex showed similar environmental susceptibility, entire morphological groups could be at risk of extinction. Importantly, if cryptic species are not recognised, assessments of ecological resilience may be biased, and substantial losses of species and genetic diversity could go undetected.

While cryptic species are often reported to occupy different environmental niches, we found no strong patterns of spatial separation among the three *S. pistillata* taxa sampled in our study (Figure 2B). We do note, however, that *S. pistillata* Taxon5 was found at the fewest sites, and at most of these sites (4 out of 5) it was found at shallower depths (Figure S8). Thus, our relatively small sample numbers might dampen our power to detect habitat preferences, and more extensive sampling (including at additional reefs), with a focus on sampling across the depth gradient, could reveal habitat specialisation.

Notably, in addition to differences in ultimate survival outcome among taxa, we also identified differences in the trajectories of bleaching and survival. While Taxon1 and Taxon5 colonies had similar survival trajectories during the early heat stress accumulation phase (i.e., before peak DHW, Figure S9), they showed different long-term outcomes, with Taxon1 exhibiting higher mortality than Taxon5 after peak DHW (Figure S9). These results imply that short-term (i.e., a few days or weeks) observations, either *in situ* or under experimental exposure, may not be reliable predictors of long-term resilience. This finding is aligned with observations of delayed impacts of heat stress in other coral species, such as increased mortality in heated *A. hyacinthus* coral fragments up to 6-month post-heat stress (Walker et al., 2022). Thus, although acute heat stress assays have become popular to test coral thermal limits (Klepac et al., 2024; Nielsen et al., 2022; Voolstra et al., 2020), observations under more realistic conditions show that responses during a recovery phase might shape the final outcome of an extreme event.

### Strong host-symbiont specificity limits separation of host and symbiont effects

Coral photosynthetic dinoflagellates vary extensively in how they are transmitted among hosts, which can influence how host-specific they may be. Here we found strong host-symbiont specificity (Figure 2C), consistent with expectations for a brooding coral species with maternal symbiont transmission (Baird et al., 2009; Bongaerts et al., 2015). Because host and symbiont taxon identity were highly correlated, we could not estimate their relative contributions to the observed thermal stress responses. However, our results appear consistent with an earlier study of *S. pistillata* corals at Heron Island Reef, which reported that colonies hosting C8 (here, exclusive to Taxon1) and C78 (here, found in all Taxon5) *Symbiodiniaceae* species bleached less than those associated with C79 and C35 (here, exclusive to Taxon4) (Sampayo et al., 2008).

Interestingly, the only surviving Taxon4 colony (both *ex situ* and on the reef) harboured C78 symbionts and one Taxon1 colony surviving both *ex situ* and on the reef switched from C8 to C78 during the common garden. C78 symbionts were associated with all colonies of Taxon5, which showed the highest survival rate, suggesting that this symbiont may provide greater thermal tolerance. Additionally, the most sensitive Taxon4 had the greatest *Symbiodiniaceae* diversity, which may reflect the re-acquisition of different symbiont taxa during past bleaching events. This shuffling could have occurred during the lifespan of the sampled colonies or in a previous generation if shuffled symbionts were inherited via vertical symbiont transmission (Quigley et al., 2019). Overall, these observations suggest that *Symbiodiniaceae* community composition may be a key contributor to response variation among *S. pistillata* holobionts, consistent with the broader role of microbiomes in host fitness (e.g., (Gould et al., 2018; Houwenhuyse et al., 2021; Petersen et al., 2023). Manipulative experiments associating each host taxon with different symbionts prior to heat stress could help clarify the relative contributions of the host and the symbionts.

### Conspecifics from historically warmer habitats show greater thermal susceptibility

Significant correlation between common garden and *in situ* survival demonstrated that intrinsic differences among coral colonies, not differential heat exposure on the reef, contributed to thermal stress response variation. However, intrinsic factors may themselves reflect historical heterogeneity in environmental experiences. Across a wide range of taxa, there is extensive evidence that organisms can be locally adapted or long-term acclimatized to micro-habitats (reviewed in Denney et al., 2020). This has also been evidenced in some coral species, where conspecifics sampled at distinct reef habitats or depths differed in their tolerance to heat stress (e.g., Bongaerts et al., 2011; Drury and Lirman, 2021; Thomas et al., 2022). Here, we investigated whether historical thermal conditions at our sampling site could explain phenotypic variation among Taxon1 colonies under common garden conditions. Whilst weak, we detected evidence for faster and more-severe bleaching for Taxon1 coral colonies originating from sites that experience greater seasonal variability and/or the warmest monthly peaks (Western windward side of the reef). These patterns also aligned with greater Taxon1 colony mortality *in situ* at Heron Bommie and Coral Garden (Figure 3A, Figure S5). However, these observations are inconsistent with local adaptation or adaptive long-term acclimatisation, where corals originating from warmer or more thermally variable habitats should show higher survival and reduced bleaching.

These counterintuitive results may arise because adaptation to historical temperature averages may not be beneficial under present-day and future acute thermal stress events. Additionally, the strong host-symbiont fidelity in this species complex limits the potential for acquiring more heat-tolerant symbionts for corals that have previously experienced bleaching, a mechanism proposed to enhance thermal tolerance in other species (Berkelmans & van Oppen, 2006). A complementary explanation is that chronic exposure to sub-lethal temperatures can induce carryover effects in the form of passive damage accumulation (Somero, 2010; Williams et al., 2016). Thus, *S. pistillata* corals from warmer habitats might be living at their upper thermal tolerance limits and have reduced capacity to cope with additional heat stress during a heatwave. Such physiological carryover effects have been documented in a range of organisms (Molina et al., 2025; Stillman et al., 2025) and previous observations of more severe bleaching in historically warmer coral reef habitats (Brown et al., 2023) are consistent with our observations. Future investigations of coral survival during heatwaves should therefore not assume that corals are locally adapted and should consider prior physiological stress. This would extend our understanding of this phenomenon and determine how it may impact our predictions of future biodiversity changes.

## Conclusion

We found that intrinsic sensitivity, but not differential exposure, was key in shaping *S. pistillata* coral thermal tolerance. Cryptic species identity and their associated symbionts strongly influenced colony-level bleaching and survival, responses under common garden conditions predicted survival on the reef and habitat thermal history may contribute further variation among individuals. Our findings highlight the importance of separating intrinsic from extrinsic factors to understand why individuals are differentially impacted during extreme climatic events. Such insights are critical to improve predictions of future losses of species and genetic diversity.

## Acknowledgments

We thank the Traditional Owners of Heron Island Sea Country, the Gooreng Gooreng, Gurang, Bailai, and Taribeland Bunda peoples, for giving their free, prior and informed consent to conduct this research as part of the Reef Restoration and Adaptation Program. We thank all the Heron Island Research Station staff members for their assistance in the field as well as M. Le Nohaïc, E. M. Sampayo, and M. A. Mello-Athayde for their advice prior to running this experiment. We also thank V. Tettamanti for assistance with symbiont typing lab work. This work was supported by the Reef Restoration and Adaptation Program, funded by the partnership between the Australian Government’s Reef Trust and the Great Barrier Reef Foundation, and by an Australian Government Research Training Program (RTP).

