## Supplementary Materials for "Heatwave winners and losers: cryptic coral holobionts differ in thermal tolerance"

**Supplementary Material for “Heatwave winners and losers: cryptic coral holobionts differ in thermal tolerance”**

**Supplementary Methods**

*Common garden experimental design*

We installed the common garden on two outdoor tables, which were shaded with neutral density dark-green clothes. On each table, we installed three sumps, connected to two aquarium tanks each. The sumps were fed seawater at a flow rate of 720L/h and included a 1500L/h pump and fed into the tanks at a rate of 144L/h. The tanks drained back into their respective sump and the sumps drained onto the holding tables. The water in the sumps was fully replaced every 4 minutes, resulting in a semi flow-through system. Fragments were randomly placed into the tanks (20 fragments per tank) ensuring that the three replicate fragments per coral colony were not in the same tank and did not share the same sump. Each tank included a 2500L/h wave maker and every other tank (one tank per sump) included a HOBO pendant MX Temp/Light logger that recorded temperature every 5 minutes.

*Whole genome* *read processing and variant calling*

Pre-processing of raw sequencing data included quality filtering, adapter removal, and alignment to a reference genome. We used Trimmomatic (Bolger et al., 2014) to remove bases with a phred-score quality less than 20 with a sliding window of 4 bp, to remove adapter sequences and to remove reads shorter than 50 bp after trimming. Trimmed paired reads were then mapped to the *S. pistillata* reference genome (GenBank assembly GCA_032172095.1) using BWA-MEM (Li & Durbin, 2009) with default settings. Resulting SAM files were converted to sorted and indexed BAM files using Samtools v1.10 (Danecek et al., 2021). We assigned read groups to BAM files and removed PCR duplicates using picard.

We followed a standard GATK pipeline to call variants (Gamer et al., 2019), using HaplotypeCaller for each individual and GenomicsDBImport to consolidate the resulting GVCF files into a Genomics Database. We then used GenotypeGVCFs to perform joint genotype calling across samples, GatherVcfs to gather variant files into a single VCF file and SelectVariants to select SNP variants only. The dataset was further filtered using VCFtools v0.1.16 (Danecek et al., 2011) with the following parameters: *--min-alleles 2, --max-alleles 2, --remove-indels, --mac 3, --minQ 30, --minGQ 20, --min-meanDP 10, --max-meanDP 20, --max-missing 0.8*.

To identify pairs of clonal samples, we performed identify by state analyses using PLINK v2.0 ([www.cog-genomics.org/plink/2.0/](http://www.cog-genomics.org/plink/2.0/), Chang et al., 2015). Then, we investigated genetic structure across samples by performing Principal Component Analysis (PCA), also using PLINK v2.0. We removed SNPs in linkage disequilibrium from the ‘all individuals’ dataset (*--indep-pairwise 200 20 0.5*) prior to running the PCA.

*Degree Heating Weeks calculation*

Degree Heating Week (DHW; °C-wk) is calculated by summing daily temperatures that exceed the locations’ climatological Maximum Monthly Mean (MMM) by 1°C (hotspots, HS) over a 12-week rolling window (Eakin et al., 2010; Liu et al., 2014):

${HS}_{i} = (T_{i} \geq MMM + 1^{\circ} C) - MMM, {HS}_{i} \geq0$

$${DHW}_{i}= \sum_{n= i -84}^{i} (\frac{{HS}_{n}}{7} ), where {HS}_{n}\geq1$$

The climatological MMM for Heron Island Reef is 27.3° C (Weeks et al., 2008).

To capture heat stress already accumulated on the reef prior to the start of the common garden experiment, we extracted regional NOAA Coral Reef Watch temperature data from the 12 weeks preceding our experiments (1 December 2023 to 3 February 2024) (NOAA CRW, 2024).

*Survival and bleaching response differences among taxa*

Taxon-specific differences in survival trajectories were tested using a mixed-effects Cox proportional hazards model. The response variable was time-to-survival for each fragment at each timepoint, with survival coded as a binary variable (alive = 1, dead = 0). The model took the following form:

coxme(Surv(Time, Survival) ~ Taxon + (1 | Taxon / Colony) + (1 | Sump / Tank))

Taxon-specific differences in phenotypic responses were tested using Generalised Mixed Linear models. Survival time, time without bleaching, and recovery time were modelled using a negative binomial distribution with log link, which accommodates overdispersion in the count data. Maximum bleaching area was modelled using a beta distribution with a logit link, which is appropriate for continuous proportion data between 0 and 1. The models took the following form:

Phenotypic trait ~ Taxon + (1 | Taxon / Colony) + (1 | Sump / Tank)

*Survival and bleaching probability trajectories along the DHW exposure*

As apparent from the ‘pyramid’ shape of the DHW curve (Figure 1C), the same DHW values occurred both as heat stress accumulate and as it dissipated. To ensure clear interpretation of taxon responses to DHW, we analysed responses before and after peak DHW separately (peak DHW = 16.1°C-wk, reached on 06/03/2024). Bleaching, data was coded as a binary outcome (unbleached or bleached) and similar to the survival model above, the models took the following form:

coxme(Surv(DHW, Survival) ~ Taxon + (1 | Taxon / Colony) + (1 | Sump / Tank))

coxme(Surv(DHW, Bleaching) ~ Taxon + (1 | Taxon / Colony) + (1 | Sump / Tank))

*Survival and bleaching response differences within taxon among environments*

Differences in phenotypic responses among conspecific coral colonies originating from different thermal environments were tested using Generalised Mixed Linear models. The models took the following form:

Phenotypic trait ~ temp-PC1 + temp-PC2 + (1 | Colony) + (1 | Sump / Tank)

**Supplementary Results**

We identified one sample that might be of hybrid origin between Taxon4 and Taxon5. In the main manuscript, we assigned this sample to Taxon4, as it showed more affinity with this group on the PCA and greater ancestry proportion with this group according to ADMITURE. However, we repeated all statistical analyses without this individual to confirm that the conclusions presented in the main text remain robust.

Specifically, survival analyses continued to support taxon-specific responses, with statistically significant survival differences among all taxa across the entire duration of the experiment (χ² = 14.1, *p* = 0.0008), between Taxon4 and the other taxa before peak DHW (χ² = 35.3, *p* = 2.08e-08) and among all taxa after peak DHW (χ² = 13.8, *p* = 0.0009).

Similarly, generalised linear mixed models continued to support significant differences among taxa in the average number of weeks they remained alive in the common garden (χ² = 38.2, *p* = 4.89e-09), in the number of weeks without bleaching (χ² = 72.2, *p* < 2e-16), and in the maximum bleaching area (χ² =59.7, *p* = 1.0e-13). As in the original analyses, recovery time did not differ significantly between Taxon1 and Taxon5 (χ² = 3.4, *p* = 0.17).

For the comparison between common garden and on-reef survival, we again found a significant association (Kappa = 0.103, z = 3.2, *p* = 0.00138) with a Spearman's correlation coefficient of ρ = 0.59. The association between common garden and on-reef survival also remained significant within each taxon (Taxon1: Kappa = 0.0871, z = 2.24, *p* = 0.0252; Taxon4: Kappa = 0.0909, z = 1.05, *p* = 0.292; Taxon5: Kappa = 0.118, z = 1.95, *p* = 0.0510).

**Supplementary Figures**

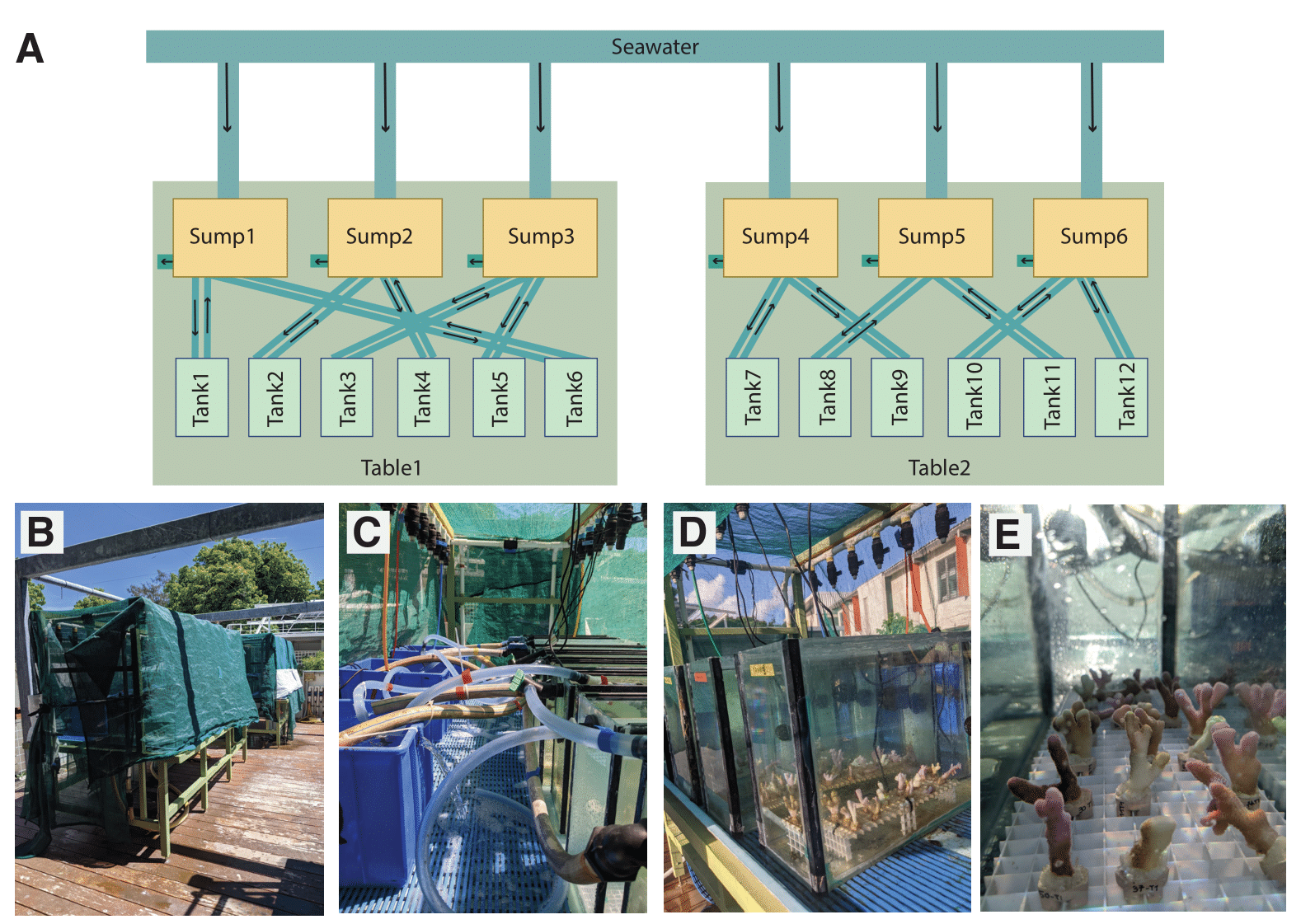

**Figure S1. Common garden experiment design and set up.** A) Experimental design diagram showing that running seawater was distributed into six sumps (Sump1–Sump6). Each sump was connected to two tanks, creating semi-flow-through water circulation systems. Sumps and tanks were organised across two separate tables (Table1 and Table2), each containing three sumps and six tanks. Photographs show how this design was executed, with the two tables with shaded clothes (B), the tubing connecting the sumps and the tanks for one table (C); the aquarium tanks lined up on one table (D) and the coral fragments placed in one of the tanks (E).

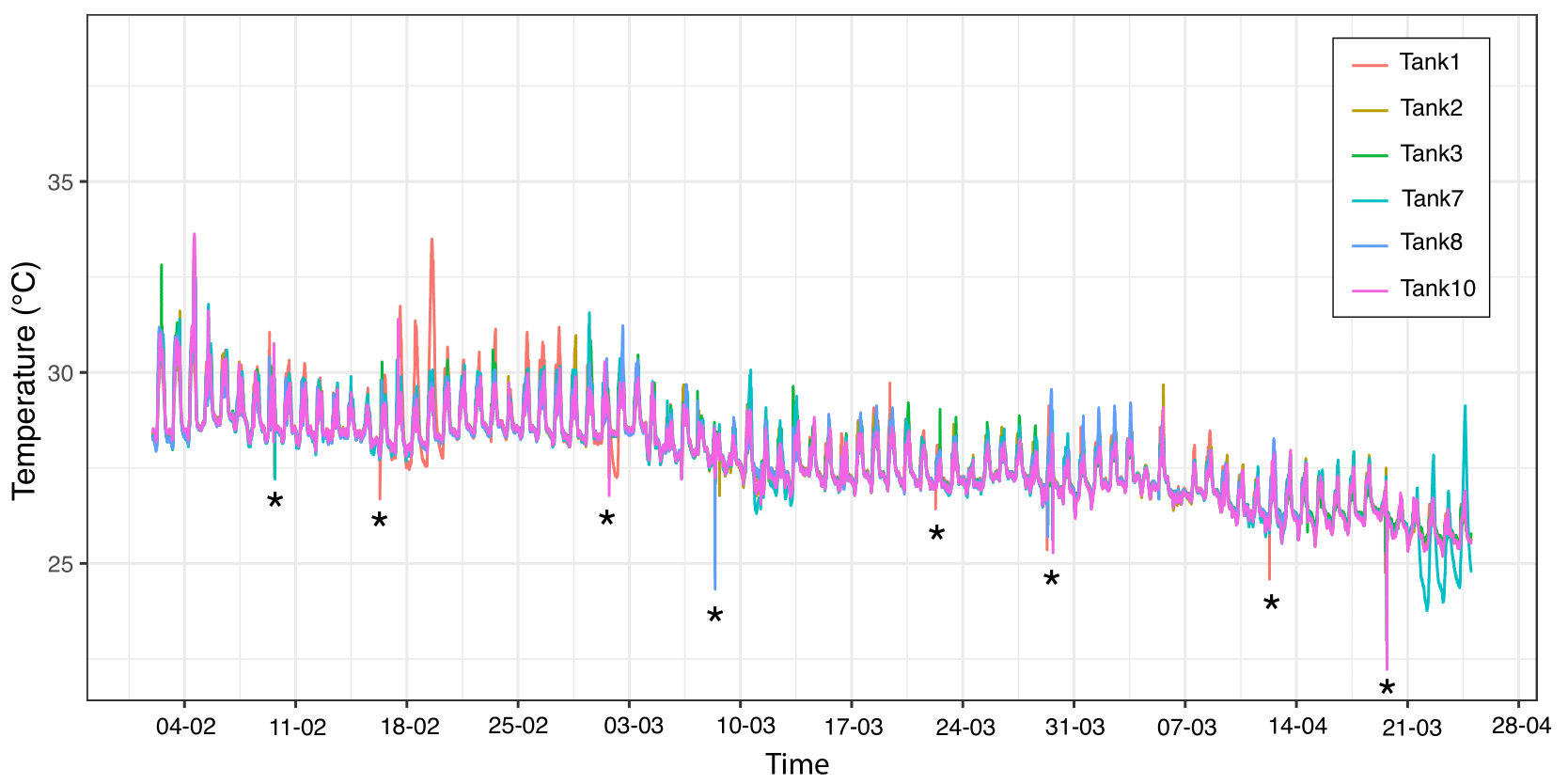

**Figure S2. Logger-recorded temperature in experimental tanks during 12 weeks of common garden experiment.** One HOBBO logger was included in every second tank (two tanks per sump, all six sumps represented, see Figure S1). Sudden drops in temperature (indicated with asterisks) correspond to periods when loggers were temporarily removed from the tanks for cleaning.

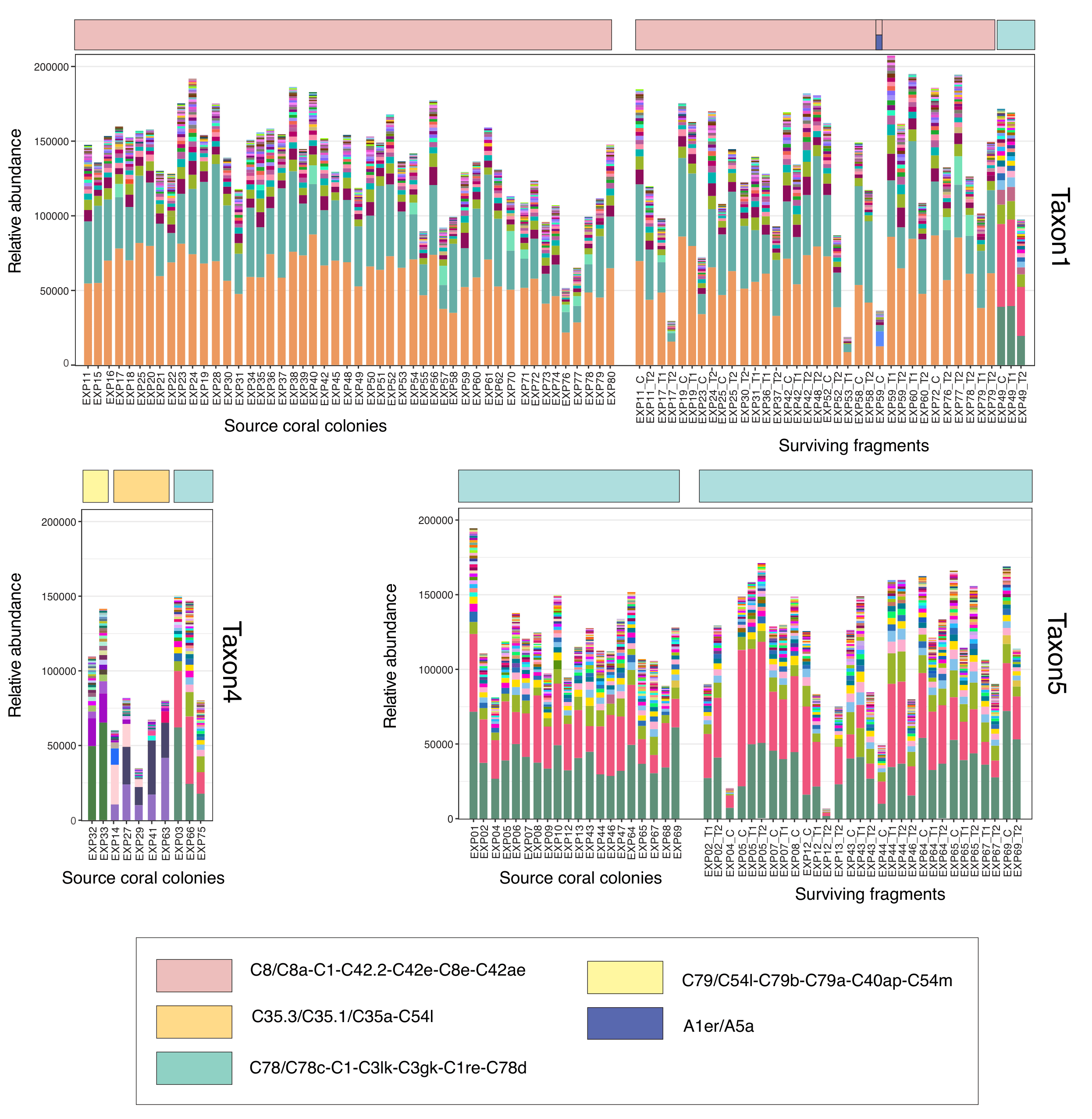

**Figure S3. Relative of ITS2 sequence variants and profiles for all samples.** For each taxon, we show relative abundance of ITS2 sequence variants of all source colonies and surviving fragments, as well as the assigned ITS2 type profile (top bar).

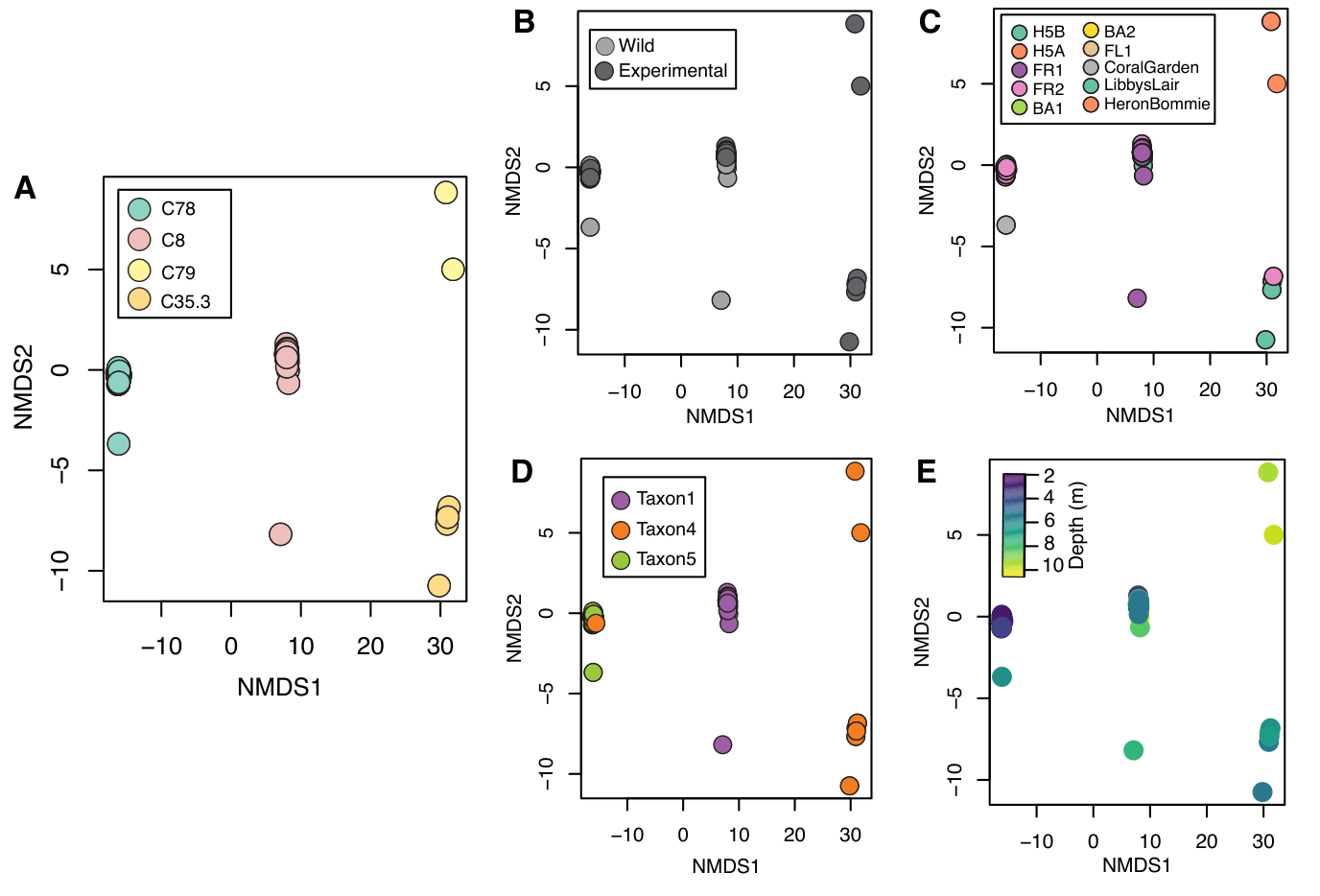

**Figure S4.** Non-metric multidimensional scaling (NMDS) ordinations based on Bray–Curtis dissimilarities of ITS2 sequence variant (DIV) abundances. Each point represents a sample, and distances between points reflect differences in symbiont community composition. Panels show the same ordination coloured by different explanatory variables: A) the ITS2 type profile as inferred from SymPortal; B) the type of sample (source wild coral colonies or fragment that survived the heat stress under common garden experimental conditions); C) sampling location;
D) coral host taxonomic identity (*S. pistillata* Taxon1, Taxon4 or Taxon5); D) depth of the source colony (in meters).

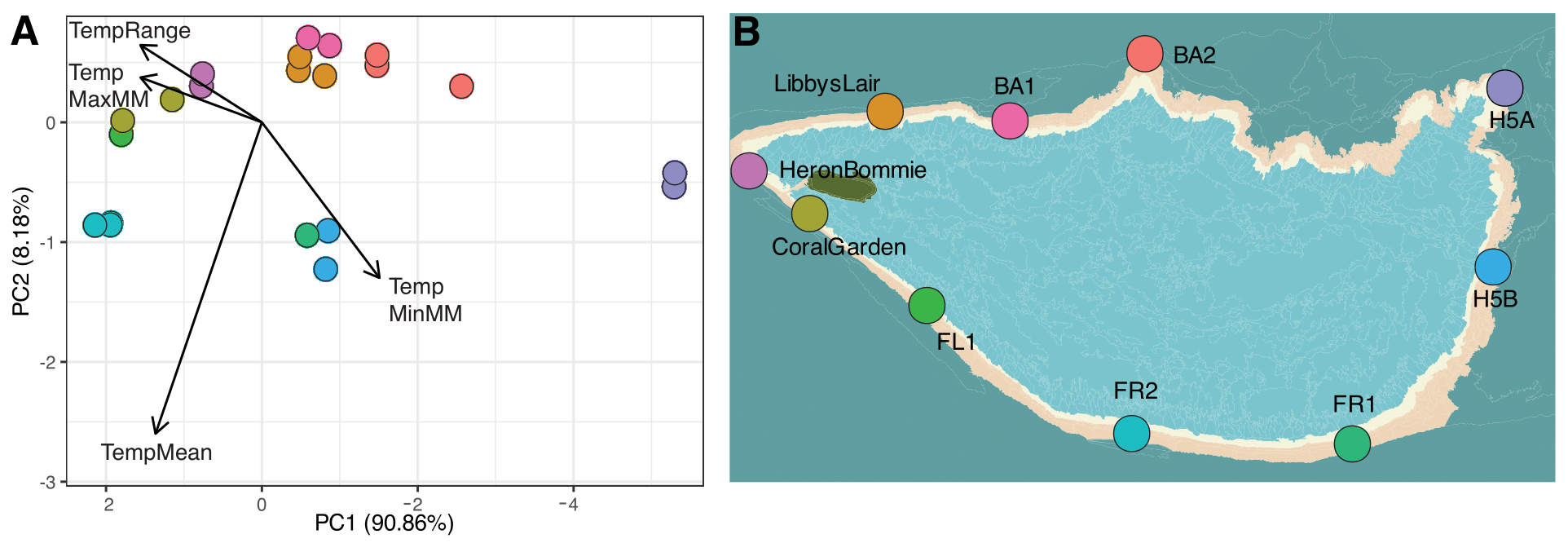

**Figure S5. Environmental PCA axes match geographic location.** A) PC1 and PC2 of a PCA summarising thermal environmental variables (mean temperature, maximum monthly mean, minimum monthly mean and temperature range; obtained via eReefs database). Due to sampling across depths, there are 21 temperature profiles in total for all sampling sites (eReefs depth bins: 0.5-2.5m, 2.5-5.25m, 5.25-9m, 9-13m); B) Map showing the geographic location of all the sampling sites. Note: PC1 direction is flipped to match geography, such that higher PC1 scores (towards the left) are associated with higher and more variable temperatures.

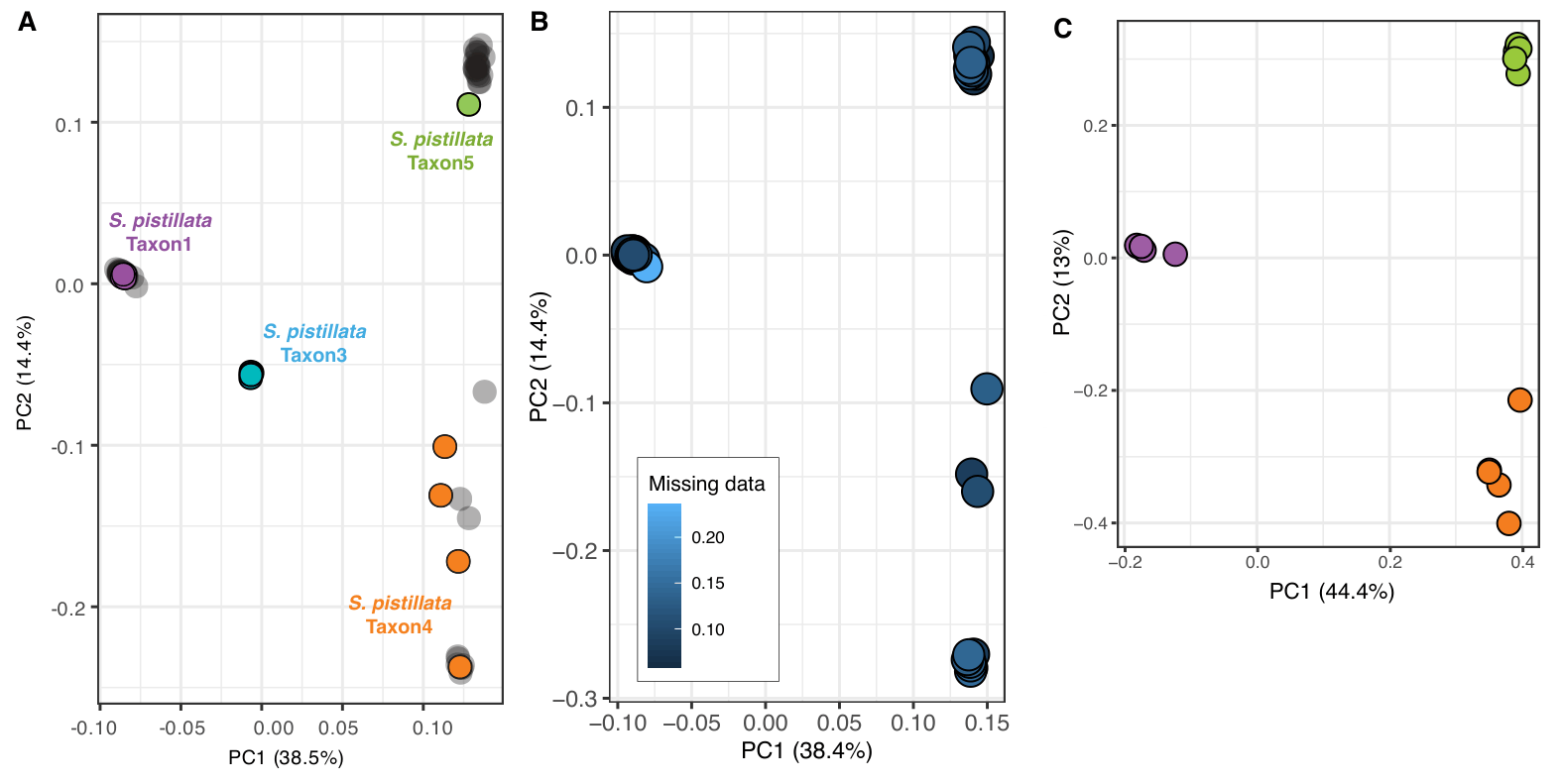

**Figure S6. Assignment of taxonomic identity for samples from this study using genomic data of representative samples from previously described *Stylophora pistillata* taxa.** (A) Principal Components Analysis based on genomic data (784,064 SNPs, 95 samples) shows the genetic similarity with samples from this study (in grey) and reference samples from previously described taxa (in block colours) to assign samples a taxonomic identity; (B) Principal Components Analysis based on genomic data (712,815 SNPs, 80 samples) shows that the intermediate samples do not have higher missing data; (C) Principal Components Analysis based on genomic data (712,815 SNPs, 15 samples) after down sampling samples from this study to 5 samples per genomic cluster, shows that the intermediate samples cluster more tightly within the Taxon4 genomic cluster.

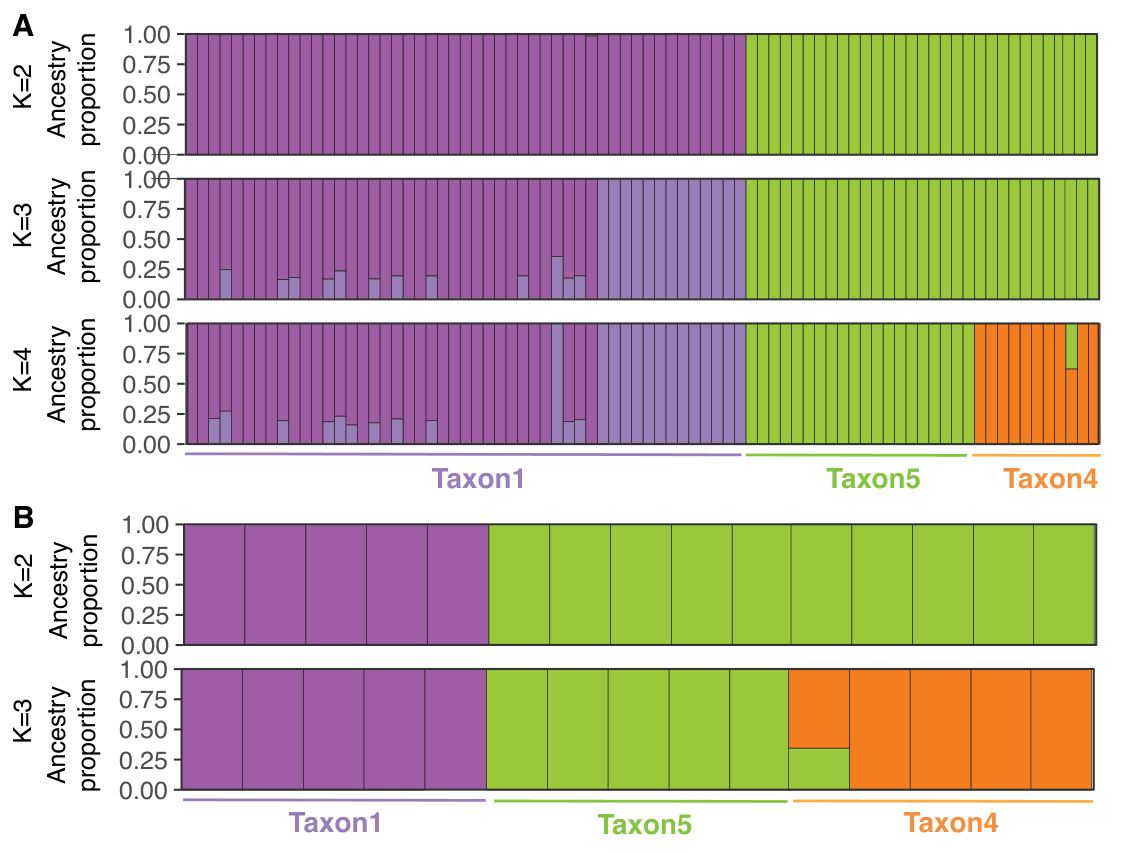
**Figure S7. Population structure inferred using ADMIXTURE across multiple K values.** (A) ADMIXTURE bar plots for all individuals, shown for K = 2–4. Each vertical bar represents an individual, and colours indicate the proportion of ancestry assigned to each genetic cluster. (B) ADMIXTURE results for a subset of five individuals per taxon, shown for K = 2–3. Colours are consistent across K values and between panels.

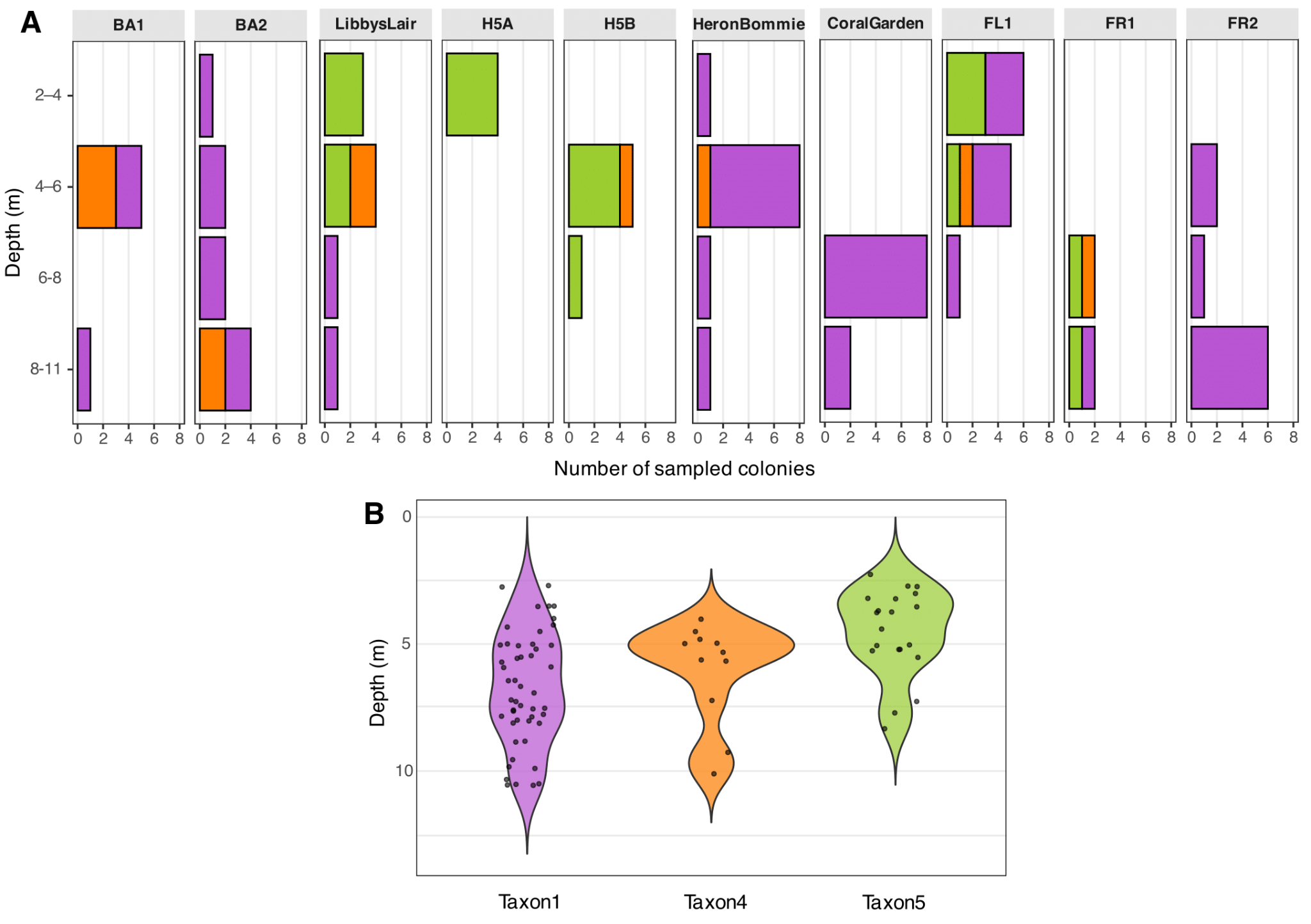

**Figure S8. *Stylophora pistillata* taxa at Heron Island Reef are sympatric across shallow depths.** A) Each panel represents one sampling site, and bars indicate the number of colonies sampled within each depth bin (0–4 m, 4–8 m, and 8–12 m). B) Depth distribution of the three taxa across all sampling sites.

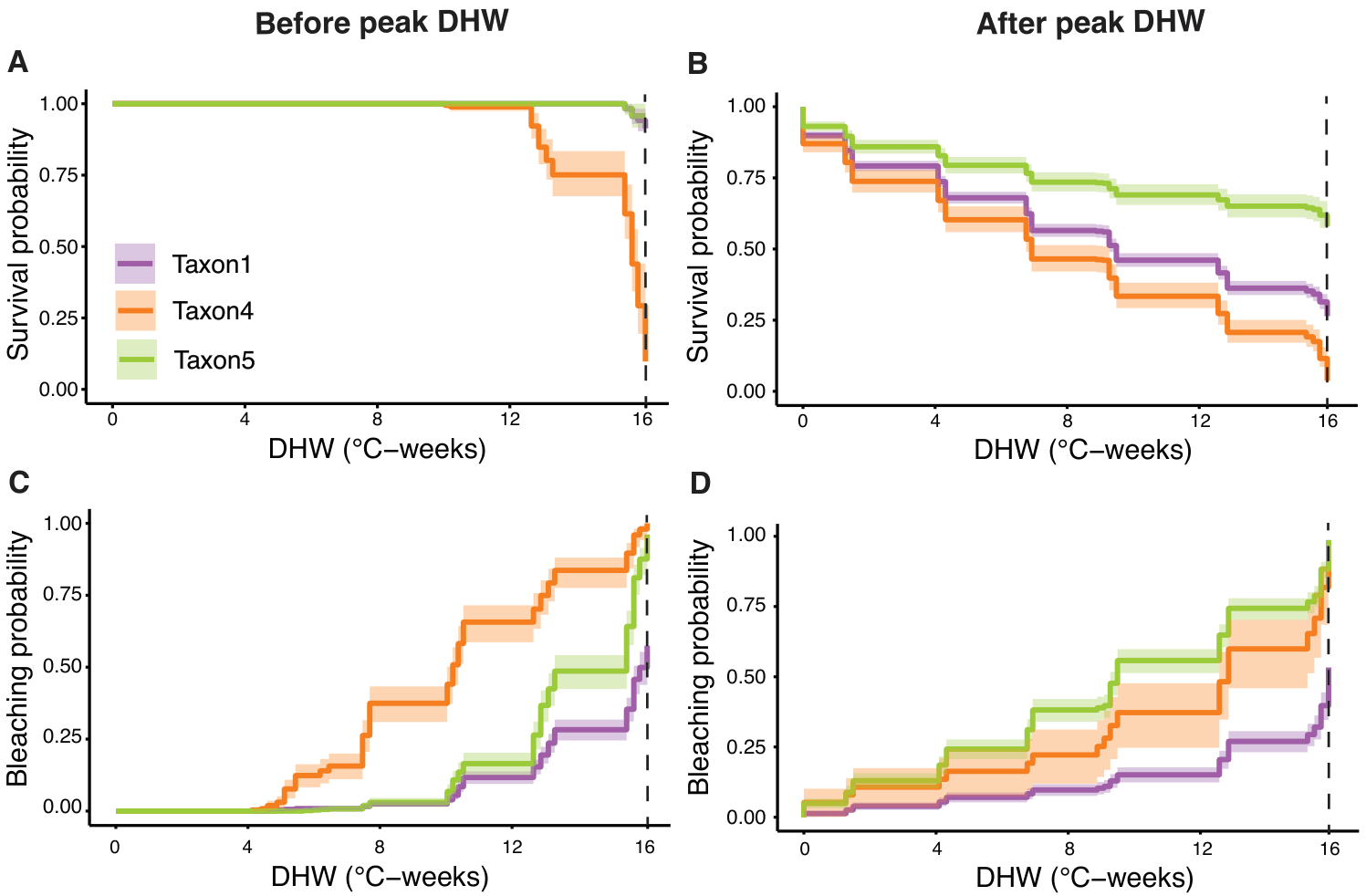

**Figure S9. Differences in** **survival and bleaching responses among *Stylophora pistillata* taxa under thermal stress, before and after peak Degree Heating Weeks (DHW).** The predicted survival and bleaching curves from mixed-effects Cox models show taxon-level differences in coral fragment survival while controlling for colony as a random effect. A) the probability of survival before peak DHW; B) the probability of survival after peak DHW; C) the probability of bleaching (full or partial) before peak DHW and D) the probability of bleaching (full or partial) after peak DHW. The dashed line represents the peak DHW. Colours used to represent the three taxa match the colours used in Figure 2. Note: Although the plots before and after peak DHW both span 0–16 DHW, their alignment with the experimental timeline differs. Panels A and C (before peak DHW) follow the forward progression of the experiment (0 DHW at the start to 16 DHW at ~5 weeks). In contrast, Panels B and D (after peak DHW) are oriented in reverse (0 DHW at the end of the experiment to 16 DHW at ~6 weeks). That is, after peak DHW, survival probability is lowest at the highest thermal stress (B), while bleaching probability is highest at the highest thermal stress (D).

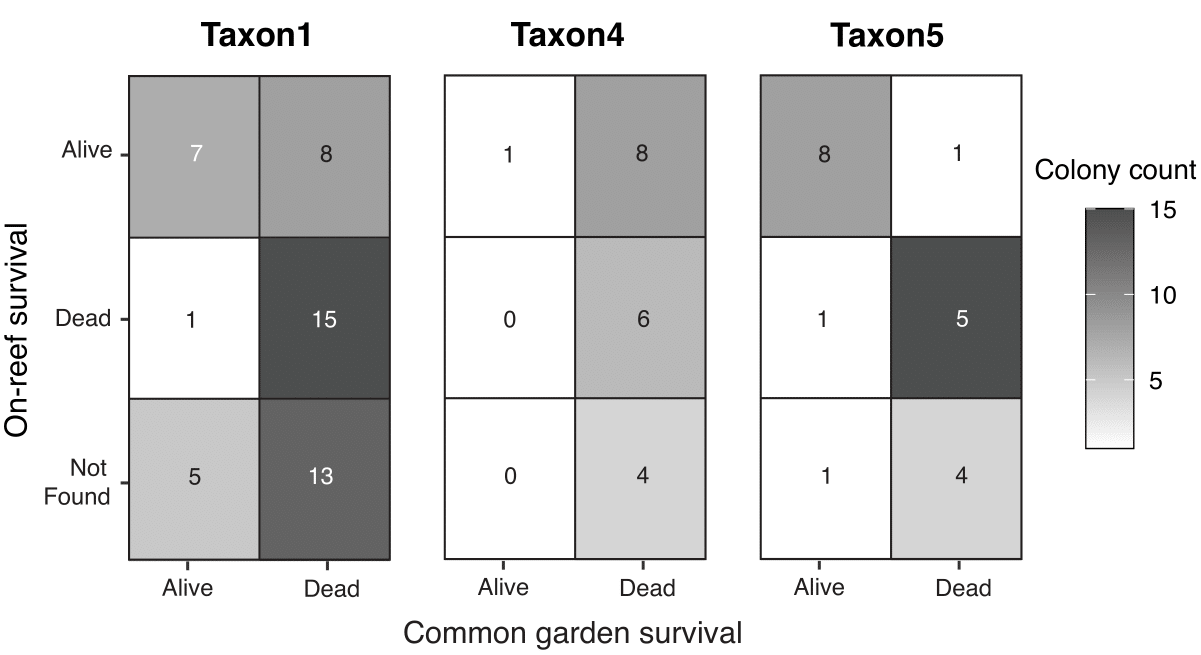

**Figure S10. Comparison of survival of *Stylophora pistillata* Taxon1, Taxon4 and Taxon5 coral colonies after the heatwave on the reef and under common garden conditions.** For each taxon, a heatmap shows the relationship between the survival of the 80 colonies on-reef following the heatwave (y-axis, rows) and in the common garden experiment (x-axis, columns). Tile colour represents the number of colonies that fall into a given combination of on-reef and common garden outcomes. Numbers inside tiles indicate the number of colonies observed for each combination (*e.g.*, for Taxon1, of the 15 colonies Alive *in situ* post-heatwave, 7 were categorised as Alive in the common garden, and 8 as Dead).

**Supplementary Tables**

**Table S1. Coordinates and historical thermal environment for each sampling site.** Monthly temperature data were extracted from the eReefs GBR1 database, across different depth categories predefined by the model. Except FR1, samples were collected from at least 2 depth bins per site, with the shallowest listed first. LL= Libby’s lair; CG= Coral Garden; HB= Heron Bommie.

| **Site** | **Lat** | | **Long** | **Depth**  **range**  **(meters)** | **Max monthly mean**  **(°C)** | **Min monthly mean**  **(°C)** | **Monthly range**  **(°C)** | **Monthly mean**  **(°C)** | **env-**  **PC1**  **score** | **env-**  **PC2**  **score** |
| --- | --- | --- | --- | --- | --- | --- | --- | --- | --- | --- |
| **H5A** | | -23.45 | 151.99 | 2.35-5.25 | 28.16 | 21.75 | 6.41 | 24.73 | -5.3 | -0.42 |
|  | |  |  | 5.25-9 | 28.16 | 21.76 | 6.41 | 24.73 | -5.28 | -0.53 |
| **H5B** | | -23.43 | 151.99 | 0.5-2.35 | 28.33 | 21.71 | 6.62 | 24.78 | -0.84 | -0.9 |
|  | |  |  | 2.35-5.25 | 28.34 | 21.72 | 6.62 | 24.79 | -0.81 | -1.22 |
| **FR1** | | -23.47 | 151.97 | 5.25-9 | 28.31 | 21.7 | 6.62 | 24.79 | -0.58 | -0.94 |
| **FR2** | | -23.47 | 151.95 | 2.35-5.25 | 28.41 | 21.66 | 6.75 | 24.82 | 2.14 | -0.85 |
|  | |  |  | 5.25-9 | 28.4 | 21.66 | 6.74 | 24.81 | -1.94 | -0.85 |
|  | |  |  | 9.0-13.0 | 28.4 | 21.66 | 6.74 | 24.81 | -1.94 | -0.84 |
| **BA1** | | -23.42 | 151.95 | 5.25-9 | 28.3 | 21.69 | 6.61 | 24.75 | -1.48 | 0.56 |
|  | |  |  | 9.0-13.0 | 28.26 | 21.71 | 6.55 | 24.74 | -1.48 | 0.47 |
| **BA2** | | -23.43 | 151.92 | 2.35-5.25 | 28.34 | 21.68 | 6.66 | 24.76 | -0.48 | 0.54 |
|  | |  |  | 5.25-9 | 28.34 | 21.69 | 6.66 | 24.76 | -0.47 | 0.43 |
|  | |  |  | 9.0-13.0 | 28.33 | 21.69 | 6.64 | 24.76 | -0.8 | 0.39 |
| **FL1** | | -23.45 | 151.92 | 2.35-5.25 | 28.4 | 21.66 | 6.75 | 24.79 | 1.79 | 0.01 |
|  | |  |  | 5.25-9 | 28.41 | 21.66 | 6.75 | 24.8 | 1.8 | -0.1 |
| **CG** | | -23.44 | 151.91 | 2.35-5.25 | 28.4 | 21.66 | 6.75 | 24.79 | 1.79 | 0.01 |
|  | |  |  | 5.25-9 | 28.39 | 21.67 | 6.73 | 24.78 | 1.15 | 0.19 |
| **LL** | | -23.43 | 151.93 | 2.35-5.25 | 28.33 | 21.68 | 6.65 | 24.75 | -0.6 | 0.7 |
|  | |  |  | 5.25-9 | 28.32 | 21.69 | 6.64 | 24.75 | -0.86 | 0.64 |
| **HB** | | -23.44 | 151.90 | 2.35-5.25 | 28.38 | 21.67 | 6.71 | 24.77 | 0.76 | 0.4 |
|  | |  |  | 5.25-9 | 28.38 | 21.67 | 6.71 | 24.78 | 0.78 | 0.3 |

**Table S2. Summary of host genotype metadata and whole-genome sequencing information.** For each sample, we report the sampling site, the sampling depth (in meters), the taxonomic identity (assigned in this study), the number of raw sequences, the **percentage of bases in the sequencing run with a Phred quality score ≥ 30 (Q30),** the sequencing depth after variant filtering and the proportion of missing data after variant filtering. Sequence data was deposited on NCBI (BioProject PRJNA1358725) and metadata was deposited on GEOME (GUID https://n2t.net/ark:/21547/Giu2).

| Sample name | Sampling  site | Sampling depth | Taxon ID | Raw reads | Q30 | Sequencing depth | Missing data |
| --- | --- | --- | --- | --- | --- | --- | --- |
| EXP01 | H5B | 7 | Taxon5 | 22,772,459 | 93.69 | 19.6385 | 0.097435 |
| EXP02 | H5B | 5 | Taxon5 | 19,595,518 | 93.13 | 17.45 | 0.125211 |
| EXP03 | H5B | 5.6 | Taxon4 | 23,785,973 | 93.27 | 20.4571 | 0.094405 |
| EXP04 | H5B | 5.3 | Taxon5 | 23,032,434 | 93.88 | 20.6319 | 0.084597 |
| EXP05 | H5B | 5 | Taxon5 | 21,935,341 | 93.65 | 19.4346 | 0.097762 |
| EXP06 | H5B | 4.8 | Taxon5 | 30,975,021 | 94.07 | 25.8282 | 0.058541 |
| EXP07 | H5A | 3.3 | Taxon5 | 24,619,068 | 93.5 | 21.6839 | 0.080029 |
| EXP08 | H5A | 2.5 | Taxon5 | 22,323,729 | 93.66 | 20.406 | 0.093644 |
| EXP09 | H5A | 2.5 | Taxon5 | 22,566,361 | 94.23 | 20.2367 | 0.089142 |
| EXP10 | H5A | 2 | Taxon5 | 27,514,649 | 93.94 | 23.9271 | 0.069516 |
| EXP11 | FR1 | 8.8 | Taxon1 | 23,862,787 | 93.63 | 20.1312 | 0.091105 |
| EXP12 | FR1 | 8.1 | Taxon5 | 24,686,140 | 93.63 | 21.9704 | 0.079521 |
| EXP13 | FR1 | 7.5 | Taxon5 | 24,121,721 | 94.21 | 21.6414 | 0.083259 |
| EXP14 | FR1 | 7.2 | Taxon4 | 24,515,152 | 93.77 | 21.1958 | 0.088326 |
| EXP15 | FR2 | 10.5 | Taxon1 | 22,495,354 | 93.64 | 18.4304 | 0.109201 |
| EXP16 | FR2 | 10.5 | Taxon1 | 24,686,301 | 93.69 | 20.988 | 0.078433 |
| EXP17 | FR2 | 10.3 | Taxon1 | 22,598,212 | 93.71 | 18.7862 | 0.097229 |
| EXP18 | FR2 | 9.8 | Taxon1 | 23,414,431 | 93.75 | 19.2809 | 0.090347 |
| EXP19 | FR2 | 9.5 | Taxon1 | 18,933,003 | 93.72 | 17.0028 | 0.115477 |
| EXP20 | FR2 | 8.1 | Taxon1 | 20,368,307 | 93.74 | 18.2044 | 0.10364 |
| EXP21 | FR2 | 6.6 | Taxon1 | 21,259,446 | 93.63 | 18.4715 | 0.099072 |
| EXP22 | FR2 | 5.9 | Taxon1 | 20,079,850 | 93.84 | 17.6481 | 0.112509 |
| EXP23 | FR2 | 4.5 | Taxon1 | 22,213,676 | 93.83 | 19.2634 | 0.093101 |
| EXP24 | BA1 | 10.5 | Taxon1 | 24,459,543 | 94.06 | 21.1267 | 0.076749 |
| EXP25 | BA1 | 5.9 | Taxon1 | 21,470,842 | 93.74 | 17.3489 | 0.114563 |
| EXP26 | BA1 | 4.5 | Taxon4 | 18,050,982 | 93.47 | 15.9277 | 0.148814 |
| EXP27 | BA1 | 5 | Taxon4 | 17,859,630 | 93.55 | 16.3014 | 0.14634 |
| EXP28 | BA1 | 5 | Taxon1 | 20,509,200 | 93.77 | 17.3569 | 0.114446 |
| EXP29 | BA1 | 5 | Taxon4 | 20,384,992 | 93.92 | 17.8407 | 0.124544 |
| EXP30 | BA2 | 10.5 | Taxon1 | 20,725,147 | 93.87 | 17.3504 | 0.114059 |
| EXP31 | BA2 | 9.9 | Taxon1 | 24,269,033 | 94.17 | 19.6851 | 0.089208 |
| EXP32 | BA2 | 10.1 | Taxon4 | 24,227,589 | 93.84 | 21.5865 | 0.086648 |
| EXP33 | BA2 | 9.3 | Taxon4 | 21,501,550 | 94.05 | 18.8128 | 0.108425 |
| EXP34 | BA2 | 6.9 | Taxon1 | 21,033,947 | 93.92 | 18.4534 | 0.111318 |
| EXP35 | BA2 | 5.4 | Taxon1 | 22,109,782 | 93.55 | 17.8208 | 0.104758 |
| EXP36 | BA2 | 6.4 | Taxon1 | 20,016,900 | 93.85 | 17.3483 | 0.119923 |
| EXP37 | BA2 | 5.5 | Taxon1 | 19,312,780 | 93.53 | 16.308 | 0.131519 |
| EXP38 | BA2 | 3.5 | Taxon1 | 23,153,878 | 93.42 | 19.9188 | 0.086414 |
| EXP39 | FL1 | 6.4 | Taxon1 | 21,617,354 | 93.57 | 18.5251 | 0.098669 |
| EXP40 | FL1 | 5.5 | Taxon1 | 20,433,530 | 94.01 | 18.1684 | 0.105198 |
| EXP41 | FL1 | 5.3 | Taxon4 | 18,679,282 | 93.64 | 16.6415 | 0.13272 |
| EXP42 | FL1 | 5 | Taxon1 | 17,017,304 | 93.76 | 15.0139 | 0.14824 |
| EXP43 | FL1 | 4.8 | Taxon5 | 17,313,772 | 93.94 | 15.9753 | 0.146586 |
| EXP44 | FL1 | 3.5 | Taxon5 | 17,774,236 | 93.57 | 16.2813 | 0.141957 |
| EXP45 | FL1 | 3.5 | Taxon1 | 20,459,518 | 93.47 | 17.2552 | 0.114183 |
| EXP46 | FL1 | 3 | Taxon5 | 19,950,743 | 94 | 18.3141 | 0.109396 |
| EXP47 | FL1 | 3.5 | Taxon5 | 20,108,915 | 93.69 | 18.4816 | 0.109232 |
| EXP48 | FL1 | 3.5 | Taxon1 | 22,022,460 | 93.1 | 18.5521 | 0.097842 |
| EXP49 | FL1 | 2.7 | Taxon1 | 20,495,702 | 93.97 | 16.9863 | 0.123513 |
| EXP50 | FL1 | 5.2 | Taxon1 | 17,660,250 | 93.94 | 15.2706 | 0.144078 |
| EXP51 | CoralGarden | 7.5 | Taxon1 | 16,423,273 | 93.46 | 14.4745 | 0.160519 |
| EXP52 | CoralGarden | 8 | Taxon1 | 18,229,828 | 93.85 | 15.883 | 0.134644 |
| EXP53 | CoralGarden | 7.6 | Taxon1 | 20,015,971 | 93.6 | 16.6301 | 0.125667 |
| EXP54 | CoralGarden | 8.1 | Taxon1 | 23,089,439 | 93.7 | 19.3913 | 0.09101 |
| EXP55 | CoralGarden | 7.8 | Taxon1 | 21,834,433 | 93.42 | 18.5762 | 0.099333 |
| EXP56 | CoralGarden | 7.8 | Taxon1 | 20,891,887 | 94.03 | 17.9241 | 0.108031 |
| EXP57 | CoralGarden | 7.6 | Taxon1 | 19,039,652 | 93.76 | 16.7985 | 0.122858 |
| EXP58 | CoralGarden | 7.4 | Taxon1 | 21,975,001 | 93.39 | 17.611 | 0.111903 |
| EXP59 | CoralGarden | 7.2 | Taxon1 | 21,140,462 | 93.85 | 18.2217 | 0.100778 |
| EXP60 | CoralGarden | 7.2 | Taxon1 | 19,742,792 | 93.36 | 17.0729 | 0.114904 |
| EXP61 | LibbysLair | 8 | Taxon1 | 16,961,845 | 93.49 | 14.7007 | 0.162958 |
| EXP62 | LibbysLair | 7.7 | Taxon1 | 23,612,213 | 93.62 | 19.2658 | 0.089546 |
| EXP63 | LibbysLair | 5.7 | Taxon4 | 19,479,980 | 94.18 | 17.1681 | 0.127574 |
| EXP64 | LibbysLair | 5 | Taxon5 | 21,697,456 | 93.8 | 18.7485 | 0.102627 |
| EXP65 | LibbysLair | 4.2 | Taxon5 | 18,056,038 | 93.9 | 15.8635 | 0.140012 |
| EXP66 | LibbysLair | 4.8 | Taxon4 | 20,727,260 | 93.94 | 17.1209 | 0.129785 |
| EXP67 | LibbysLair | 3.5 | Taxon5 | 19,677,339 | 93.43 | 17.4391 | 0.114185 |
| EXP68 | LibbysLair | 3 | Taxon5 | 18,409,830 | 93.7 | 16.8333 | 0.1296 |
| EXP69 | LibbysLair | 2.8 | Taxon5 | 17,560,250 | 93.18 | 16.2202 | 0.135922 |
| EXP70 | HeronBommie | 8.8 | Taxon1 | 20,353,214 | 93.88 | 17.4921 | 0.112499 |
| EXP71 | HeronBommie | 7.5 | Taxon1 | 18,768,048 | 93.62 | 15.985 | 0.134555 |
| EXP72 | HeronBommie | 5.7 | Taxon1 | 24,436,018 | 93.61 | 20.5866 | 0.081905 |
| EXP73 | HeronBommie | 2.7 | Taxon1 | 20,534,665 | 94.01 | 12.7094 | 0.236397 |
| EXP74 | HeronBommie | 4 | Taxon1 | 16,402,973 | 93.43 | 14.2705 | 0.169071 |
| EXP75 | HeronBommie | 4 | Taxon4 | 18,922,025 | 93.97 | 16.4288 | 0.143859 |
| EXP76 | HeronBommie | 4.3 | Taxon1 | 17,233,196 | 93.42 | 14.9882 | 0.152812 |
| EXP77 | HeronBommie | 4.2 | Taxon1 | 17,222,430 | 93.46 | 14.7946 | 0.157007 |
| EXP78 | HeronBommie | 5 | Taxon1 | 20,475,693 | 93.44 | 17.664 | 0.109939 |
| EXP79 | HeronBommie | 5 | Taxon1 | 21,956,485 | 93.64 | 17.8649 | 0.106385 |
| EXP80 | HeronBommie | 5 | Taxon1 | 21,010,061 | 93.1 | 18.1245 | 0.106772 |

**Table S3.** **Colony-level** **estimated marginal means for four thermal stress response traits among** ***Stylophora pistillata* taxa during the common garden experiment.** Survival time, time without bleaching and recovery time are expressed in number of weeks, maximum bleaching area is expressed in percentage of total fragment area. Recovery time could not be estimated for Taxon4; 95% CI are shown in parentheses.

| **Taxon** | **Survival time** | **Time without bleaching** | **Maximum bleaching area** | **Recovery time** |
| --- | --- | --- | --- | --- |
| **Taxon 1** | 7.7 (7.0-8.5) | 5.0 (3.4-7.6) | 30 (20-40) | 2.6 (1.6-4.5) |
| **Taxon 4** | 4.6 (3.9-5.6) | 0.1 (0.08-0.4) | 87 (75-94) | - |
| **Taxon 5** | 9.7 (8.4-11.1) | 2.6 (1.5-4.8) | 78 (65-84) | 1.5 (0.9-2.6) |

**Table S4. Variance components from the generalised linear mixed models.** Summary statistics are presented for 1) the number of weeks the fragments were alive (*i.e.,* survival time); 2) the number of weeks the fragments did not show signs of bleaching (*i.e.*, time without bleaching); 3) the maximum bleached area per fragment and 4) the number of weeks it took fragments’ health scores to go up after reaching their lowest value (*i.e.*, recovery time). We report the estimates, standard errors, z-values, and p-values for the fixed effect, Taxon. Random effects were colony, sumps and tanks nested within sumps and we report their variance and standard deviations. Differences among replicate fragments per colony that are not explained by other sources (taxon, colony, sump or tank effects) are captured in the models’ residual. For each model, the estimated effect of each taxon identity is reported relative to Taxon1, along with the standard error of the estimate, the z-value (estimate divided by its standard error). The associated *p*-value tests the null hypothesis that the effect equals zero. For Taxon1, the null hypothesis is that survival time differs from zero; for Taxon4 and Taxon5, the null hypothesis is that their survival times differ from Taxon1. The marginal means from these models for each Taxon, back on the original trait measurement scale, are reported in Table 1 of the main document.

| **Formula:** Survival time ~ Taxon + (1 \| Colony) + (1 \| Sump/Tank) | | | | |
| --- | --- | --- | --- | --- |
| **Random effect** | **Variance** | **Standard deviation** | | |
| **Colony** | 8.67e-02 | 0.29 | | |
| **Tank:Sump** | 6.97e-03 | 0.083 | | |
| **Sump** | 2.44e-08 | 0.00016 | | |
| **Fixed effects** | **Estimate** | **Standard error** | **z value** | **Pr(>\|z\|)** |
| **Taxon1** | 1.95 | 0.047 | 41.85 | < 2e-16 |
| **Taxon4** | 0.092 | 0.047 | 1.97 | 0.049 |
| **Taxon5** | -0.42 | 0.067 | -6.19 | 6.17e-10 |

| **Formula:** Time without bleaching ~ Taxon + (1 \| Colony) + (1 \| Sump/Tank) | | | | |
| --- | --- | --- | --- | --- |
| **Random effect** | **Variance** | **Standard deviation** | | |
| **Colony** | 1.16e+00 | 1.076 | | |
| **Tank:Sump** | 2.32e-01 | 0.48 | | |
| **Sump** | 4.45e-06 | 0.0021 | | |
| **Fixed effects** | **Estimate** | **Standard error** | **z value** | **Pr(>\|z\|)** |
| **Taxon1** | 0.291 | 0.205 | 1.42 | 0.156 |
| **Taxon4** | 1.3439 | 0.1817 | 7.379 | 1.59e-13 |
| **Taxon5** | -2.0737 | 0.2707 | -7.470 | 8.01e-14 |

| **Formula:** Max bleaching area ~ Taxon + (1 \| Colony) + (1 \| Sump/Tank) | | | | |
| --- | --- | --- | --- | --- |
| **Random effect** | **Variance** | **Standard deviation** | | |
| **Colony** | 1.66 | 1.29 | | |
| **Tank:Sump** | 0.19 | 0.43 | | |
| **Sump** | 0.07067 | 0.27 | | |
| **Fixed effects** | **Estimate** | **Standard error** | **z value** | **Pr(>\|z\|)** |
| **Taxon1** | 0.78 | 0.24 | 3.27 | 0.0011 |
| **Taxon4** | -1.64 | 0.21 | -7.97 | 1.64e-15 |
| **Taxon5** | 1.16 | 0.29 | 4.04 | 5.29e-05 |

| **Formula:** Recovery time ~ Taxon + (1 \| Colony) + (1 \| Sump/Tank) | | | | |
| --- | --- | --- | --- | --- |
| **Random effect** | **Variance** | **Standard deviation** | | |
| **Colony** | 5.99e-01 | 0.77 | | |
| **Tank:Sump** | 3.314e-01 | 0.57 | | |
| **Sump** | 2.50e-07 | 0.00050 | | |
| **Fixed effects** | **Estimate** | **Standard error** | **z value** | **Pr(>\|z\|)** |
| **Taxon1** | 0.77 | 0.22 | 3.44 | 0.00059 |
| **Taxon4** | 0.19 | 0.18 | 1.083 | 0.29 |
| **Taxon5** | -0.0068 | 0.28 | -0.024 | 0.98 |
